# Kidney collecting duct cells make vasopressin in response to NaCl induced hypertonicity

**DOI:** 10.1101/2022.05.13.491898

**Authors:** Juan Pablo Arroyo, Andrew S. Terker, Yvonne Zuchowski, Jason A. Watts, Fabian Bock, Cameron Meyer, Wentian Luo, Meghan Kapp, Edward R. Gould, Adam X. Miranda, Joshua Carty, Ming Jiang, Roberto Vanacore, Elizabeth Hammock, Matthew H. Wilson, Roy Zent, Mingzhi Zhang, Gautam Bhave, Raymond C. Harris

## Abstract

Vasopressin has traditionally been thought to be produced by the neurohypophyseal system and then released into the circulation where it regulates water homeostasis. The syndrome of inappropriate secretion of anti-diuretic hormone (vasopressin) raised the question if vasopressin could be produced outside of the brain and whether the kidney could be a source of vasopressin. We found that mouse and human kidneys expressed vasopressin mRNA. Using an antibody that detects the pre-pro-vasopressin, we found that immunoreactive pre-pro-vasopressin protein is found in mouse and human kidneys. Moreover, we found that murine collecting duct cells make biologically active vasopressin which increases in response to NaCl mediated hypertonicity, and that water restriction increases the abundance of kidney-derived vasopressin mRNA and protein expression in mouse kidneys. Thus, we provide evidence of biologically active production of kidney-derived vasopressin in kidney tubular epithelial cells.

## Introduction

Vasopressin is a nine amino-acid peptide hormone that plays a key role in water and blood pressure homeostasis (1). Vasopressin is the end-product of a highly processed 164 amino acid pre-pro-peptide. Processing of the vasopressin pre-pro-peptide results in three distinct peptides with a 1:1:1 ratio: vasopressin, neurophysin 2, and copeptin. Vasopressin is the biologically active hormone, neurophysin 2 is a carrier protein for vasopressin, and copeptin is the c-terminal glycosylated end-product. Physiologic vasopressin production is currently thought to be limited to the brain under physiologic conditions. The main physiologic stimuli for vasopressin production in the hypothalamus are increased extra-cellular fluid tonicity and hypotension (1). Vasopressin binds to three distinct G-coupled protein receptors, V1a, V1b, and V2. The V2 receptor (V2R) is mainly expressed in the kidney along the distal nephron and is a critical regulator of mRNA transcription, protein abundance, and trafficking of the water channel, Aquaporin-2 (AQP2) (2-4). Vasopressin signaling through V2R leads to AQP2 phosphorylation and translocation to the apical membrane of connecting and collecting duct cells (5), which leads to increased apical water permeability with consequent water retention and increased urine concentration.

Unregulated vasopressin production can lead to excessive water retention and decreased serum sodium (Na^+^) levels. This common clinical scenario, known as Syndrome of Inappropriate Antidiuretic Hormone secretion (SIADH) occurs in several diseases processes including, malignancy, pulmonary disorders, central nervous system disorders, non-hypothalamic tumors, and certain medications (6). Moreover, there is a large body of work that shows that vasopressin is involved in a broad range of physiologic and pathophysiologic states that go beyond water and blood pressure homeostasis (7-20). In fact, there are multiple reports of local vasopressin mRNA synthesis outside of the brain (21-29). This raises the question of whether biologically active vasopressin is also produced outside of the brain under physiologic conditions and if this locally produced vasopressin functions independent of pathways linked to water transport and body fluid balance.

To answer this question, we investigated whether kidney epithelial cells produce vasopressin and if this production is regulated by changes in extracellular fluid tonicity. We found evidence of vasopressin gene expression and protein production in mouse and human kidney epithelial cells. We demonstrated that this vasopressin activated V2R in vitro, and its production was increased when cells were placed in hypertonic NaCl solution. Finally, we provide evidence that whole kidney vasopressin mRNA and protein expression increased in water deprived mice. Thus, we conclude that kidney epithelial cells produce vasopressin that can be increased by NaCl mediated hypertonicity under physiological conditions.

## Methods

### In-silico database analysis

To analyze expression of the AVP gene in a systematic manner, we used Nephroseq v5.0 an online platform hosted by the University of Michigan that allows integrative data mining of publicly available, peer-reviewed, genotype/phenotype data (www.nephroseq.org) (30). We then used the gene name: “AVP” and set the following thresholds: p Value 0.05, r Value 0.5, and Fold change of 1.5. Complete results of the analysis are reported in Supplementary Table 1 and 2.

### Cells

*Wild-type* C57BL/6J inner medullary collecting duct cells (IMCDs) were isolated and transfected with SV40 large T-cell antigen as described previously (31). Cells were cultured in DMEM high glucose, pyruvate containing medium supplemented with 10% FBS and 100 I.U./mL penicillin + 100 (μg/mL) streptomycin and grown to confluency on permeable supports. Cells from passages 2-7 were plated at a density of 5x10^5^ cells per well (12-well dish) and serum was withdrawn 24 hrs prior to experiments. We confirmed that our IMCDs have a gene expression profile similar to that of native IMCDs including *Aqp2*, *Aqp3*, *Aqp4*, *Slc14a2*, and *Avpr2* (Supplemental Figure 2A).

#### Hypertonic stimulation

Confluent IMCDs were serum starved for 24 hrs, after which new FBS-free DMEM was supplemented with 100 mmol NaCl in the presence or absence of decreasing doses of V2r inhibitor, OPC 31260. Cells were kept in the NaCl+/- OPC supplemented medium for 24 hrs after which cells were harvested in Trizol (Invitrogen) for RNA extraction or RIPA (Thermo Scientific) for total protein extraction. To compare hypertonic stimuli, FBS-free DMEM supplemented with 100 mmol NaCl, 200 mmol glucose, or 200 mmol mannitol was added to serum starved confluent IMCDs for 12 or 24 hrs. Cells were then harvested in Trizol (Invitrogen) for RNA extraction.

#### Conditioned medium

Confluent IMCDs were serum starved for 24 hrs, after which the control group was treated with FBS-free DMEM, and the NaCl group was treated with FBS-free DMEM supplemented with 100 mmol NaCl. After 24 hrs the medium was collected from both NaCl treated and control groups and the NaCl conditioned medium was diluted to achieve a calculated osmolality similar to that of the control medium (∼320 mOsm) (Figure 4A). The control and NaCl conditioned media were then added to confluent serum starved HEK-hV2R-CRE-Luc cells or IMCDs or (see below) for 3 hrs. The 3 hr time point was chosen since prior work by Hasler et al. showed that 3 hrs of hypertonic stimulation decreased AQP2 expression (32). IMCDs were then harvested in RIPA buffer (Thermo Scientific) or fixed on a slide with 4% PFA for immunolabeling and HEK-hV2R-CRE-Luc cells underwent the luciferase reporter assay as described below.

#### V2R luciferase reporter cell assay

*piggyBac* transposon vectors (33) were designed and ordered from Vectorbuilder (Chicago, IL) to express the human V2R along with puromycin resistance (PB-Puro-CMV-hAVPR2) or cyclic AMP (cAMP) response element driving luciferase (pPB-CREminiP-Luc) in separate vectors. HEK293 cells were transfected in 6 well plates at 60% confluence using lipofectamine LTX (Invitrogen) with 1µg of each transposon vector along with 0.5µg of pCMV-m7PB hyperactive transposase (34), according to the manufacturer’s instructions. One day after transfection, cells were split to 100mm dishes and selected with 3µg/ml of puromycin for two weeks. For luciferase assays, stably transfected cells (HEK-hV2R-CRE-Luc) were treated with different reagents as indicated for 3-18 hours in 24 well plates. Two µl of 30mg/ml of XenoLight D-Luciferin (PerkinElmer) was added to wells after treatment and incubated for 5 minutes. Luciferase expression was quantitated by capturing photons/sec using a PerkinElmer IVIS imaging system.

#### HEK-293 transient transfection

HEK-293 cells were cultured in 6 well plate to 60% confluence and transfected using lipofectamine LTX (Invitrogen) using 2.5 ug of pCMV6-XL5 or pCMV6-XL5-*AVP* (Origene) following the manufacturer’s instructions. Cells were collected in RIPA buffer (Thermo Scientific) 24 hrs after transfection.

#### siRNA transfection

IMCDs were plated at a density 7.5 x10^5^ cells per well and upon reaching 30% confluency they were transfected with Silencer Select (ThermoFisher) *Avp* siRNA (AssayID: s232140) or ctrl siRNA (AssayID: 4390843) according to the manufacturer’s instructions. Cells were lysed in RIPA buffer (Thermo Scientific) upon reaching 90% confluency 24-36 hrs later and pre-pro-vasopressin protein amount was evaluated with immunoblotting (see below).

### Mice

All animal experiments were performed in accordance with the guidelines and with the approval of the Institutional Animal Care and Use Committee of Vanderbilt University Medical Center.

*Avp^tm1.1(cre)Hze^*; *Gt(ROSA)26Sor^tm4(ACTB-tdTomato,-EGFP)Luo^* Transgenic mouse <u>– Heterozygote</u> B6.Cg-*Avp^tm1.1(cre)Hze^*/J (AVP-IRES2-Cre-D) (Jax Strain No. 023530) in which Cre-recombinase is linked to arginine vasopressin gene expression (35) was crossed with homozygous B6.129(Cg)-*Gt(ROSA)26Sor^tm4(ACTB-tdTomato,-EGFP)Luo^*/J (ROSA^mT/mG^) (Jax Strain No. 007676) a two-color fluorescent Cre-reporter allele that expresses membrane GFP in Cre-expressing cells (36) to generate the AVP-IRES2-Cre-D;ROSA^mT/mG^ reporter mouse. Age-matched, 8-week old, AVP-IRES2-Cre positive and negative, ROSAmT/mG positive mice were used. Genotype was confirmed pre and post euthanasia with Transnetyx automated genotyping. Following euthanasia, kidneys were removed and incubated at room temperature overnight in 3.7% formaldehyde, 10 mM sodium m-periodate, 40 mM phosphate buffer, and 1% acetic acid. The fixed kidneys were dehydrated through a graded series of ethanol, embedded in paraffin, sectioned (5 µm), and mounted on glass slides. Slides were then probed with primary antibodies against GFP (Aves Labs – Cat No. GFP-1020), goat anti-chicken – AF-488 secondary (Thermo – Cat. No. A32931), and co-stained with a mouse anti-AQP2-AF647 conjugated antibody (Santa Cruz - sc-515770) to identify collecting duct cells (Supplemental Figure 1).

#### Water Loading and Restriction

Age-matched male wild type C57BL/6J (8-12 weeks old) were used for the experiments. *Water restriction* - Twenty-four hours prior to euthanasia, water was removed from cages of the water restricted group. *Water loading* – Twenty-four hours prior to euthanasia, food in the water loaded group was changed to a gelled diet (37). Briefly, 65 grams of crushed 4.5% fat mouse chow (5L0D - LabDiets) was mixed with 7 g of gelatin and dissolved in 120 mL of water. Gel was solidified in plastic cups and then served as the sole source of food for 24 hrs, 9:00 AM to 9:00 AM. Ad-lib access to water was maintained throughout.

### Reverse Transcription and Real Time qPCR

RNA from cells and kidneys was isolated with Trizol reagent (Invitrogen) following the manufacturer’s instructions. cDNA was synthesized from equal amounts of total RNA from each sample using SuperScript IV First-strand Synthesis Sytem kit (Invitrogen). *Reverse Transcription* - PCR was performed using Q5 High-Fidelity DNA Polymerase (New England BioLabs), forward primer (ATGCTCGCCAGGATGCTCAACACTACG) reverse primer (TCAGTAGACCCGGGGCTTGGCAGAATCCACGGACTC). *Quantitative RT-qPCR* -was carried out using TaqMan real-time PCR (7900HT, Applied Biosystems). All gene probes and master mix were purchased from Applied Biosystems. The probes used in the experiments were: mouse *Avp* (Mm00437761_g1) (**Figure 1**) and (Mm01271704_m1) (**Supplemental Figure 2 D**), mouse *Aqp2* (Mm00437575_m1), mouse *Aqp3* (Mm01208559_m1), mouse *Aqp4* (Mm00802131_m1), mouse *Akr1b3* (Mm01135578_g1), mouse *Slc14a2* (Mm01261839_m1), mouse *Avpr2* (Mm01193534_g1), mouse RPS18 (Mm02601777).

**Figure 1.**
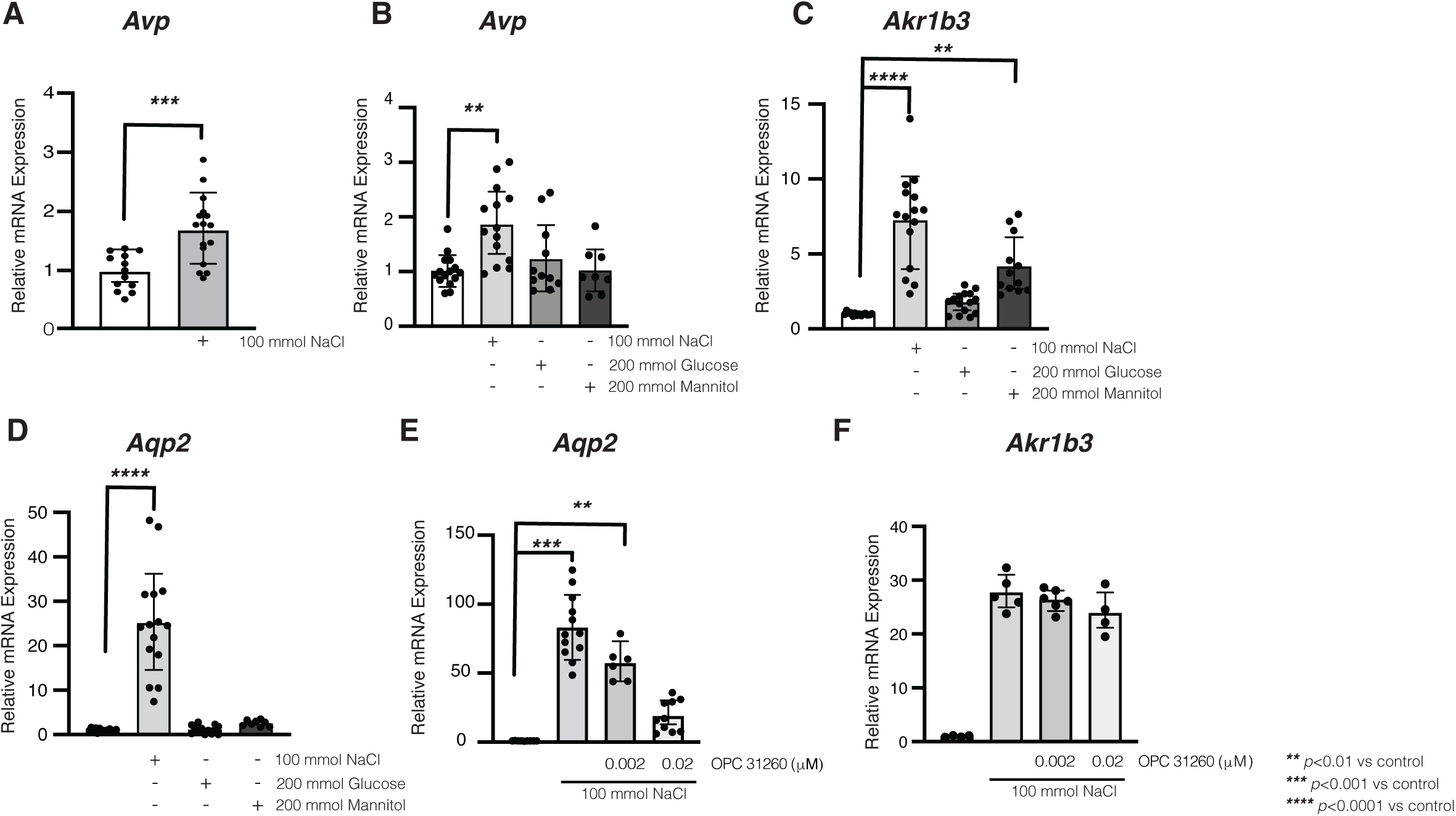
NaCl mediated hypertonicity increased vasopressin (*Avp*) mRNA in kidney epithelial cells. Incubation with NaCl (500 mOsm) for 24 hrs increased vasopressin mRNA in mouse inner medullary collecting duct cells (p<0.001 vs control) **(A)**. NaCl (p<0.01) but not mannitol or glucose increase *Avp,* mRNA **(B)**. Aldose reductase (*Akr1b3*), a marker of hypertonic stress, increased with NaCl (p<0.0001 vs control) and mannitol (p<0.001 vs control) but not glucose **(C)**. Aquaporin 2 (*Aqp2*) mRNA increased after treatment with NaCl but not glucose or mannitol **(D)**. Vasopressin receptor 2 antagonist OPC-31260 (20 nm) blunted the NaCl mediated increase in *Aqp2* (**E**) despite similar hypertonic stress (**F**). Data are presented as mean ± SD and analyzed with an unpaired Mann-Whitney (A) or 1-way ANOVA with Tukey’s post hoc analysis (B-F), with a minimum of 4 independent replicates per group ** p<0.01, *** p<0.001, and ****p<0.0001 vs control.

Droplet Digital PCR (ddPCR) – All ddPCR experiments were performed using a QX200 AutoDG Droplet Digital PCR System (Bio-Rad) with assistance from the Vanderbilt University Medical Center immunogenomics, Microbial Genetics and Single Cell Technologies Core. For all experiments plates of droplets of PCR mixture were automatically generated with an AutoDG, TaqMan probes were added, the plate was heat-sealed, and amplified with a C1000 Touch Thermal Cycler. Droplets were then read with a Droplet Reader (Bio-Rad). DNA concentrations were determined only from wells with >12,000 droplets using QuantaSoft v1.7.4.0917 after manually setting the positive droplet threshold above the signal from no-template controls in the same plate. Counts are reported as copies per 20 uL normalized to Tert gene expression. The following probes were used: mouse *Avp* (Mm00437761_g1), human *AVP* (Hs00356994_g1), TaqMan Copy number reference assay, mouse and human, Tert Cat No. 4458368, and 4403316.

RNA in situ hybridization – Experiments were performed with RNAScope reagents (ACD, Biotechne) and according to manufacturer’s instructions. Briefly, FFPE kidney sections (mouse and human) were stained using the RNAScope Multiplex Fluorescent Reagent v2 kit RED. Probes against mouse vasopressin (Mm-*Avp*) (Cat. No. 472261) mouse aquaporin 2 (Mm-*Aqp2*) (452411), human vasopressin (Hs-*AVP*) (Cat. No. 401361), and human aquaporin 2 (Hs-*AQP2*) (Cat. No. 434861) were used, the 3-Plex negative control probe provided by the manufacturer was used as a negative control.

### Antibody development

Antibody was developed with assistance from Phosphosolutions Inc. Briefly, rabbits were immunized with a synthetic peptide that corresponds to amino acid residues that span the cleavage sites between vasopressin and neurophysin 2 which were conjugated to keyhole limpet hemocyanin (KLH). Antibodies were then purified with affinity purification.

### Immunoblotting

Protein was extracted from cells and whole kidneys using RIPA buffer (Thermo Scientific) with protease and phosphatase inhibitors (Roche), total protein was then quantified with BCA Assay (Pierce), and equal amounts of protein were loaded in MiniProteanTGX polyacrylamide precast gels (Bio-Rad) and transferred to nitrocellulose using Transblot Turbo (Bio-Rad). Nitrocellulose membranes were stained for total protein using Ponceau-S (Thermo Fisher) and blocked with 5% non-fat dry milk for 1 hour room temperature. Primary antibodies used were: anti-pre-pro-vasopressin (PhosphoSolutions) 1:1000 overnight, anti-aquaporin E2 sc-515770 (Santa Cruz) 1:100 overnight, anti-aquaporin phosphor serine269 (38) (Phosphosolutions - p112-269), 1:1000 overnight. Secondary antibodies were HRP coupled anti-mouse (AB_10015289) and anti-rabbit (AB_2337938) from Jackson Immuno-Research. Band density quantification was performed with ImageJ.

### Immunoprecipitation

Magnetic protein G beads (Sigma) were used per manufacturer’s instructions. Briefly, protein G beads were coupled to the pre-pro-vasopressin antibody or normal rabbit serum (Sigma) at a 1:50 ratio for 10 min at room temperature, after which one milligram of IMCD whole cell lysate was added to the antibody coupled beads and incubated for 1 hr room temperature. Proteins were then eluted, mixed with 2X Laemmli buffer (Bio-Rad) and run on polyacrilamyide gels, transferred to nitrocellulose membranes and incubated with anti-pre-pro-vasopressin (PhosphoSolutions) 1:1000 1 hr room temperature, anti-vasopressin (AB 1565 Millipore Sigma) 1:1000 (39, 40) 1 hr room temperature, or anti-neurophysin 2 (MABN856 – Sigma) (41) followed by HRP coupled anti-rabbit (AB_2337938) or anti-mouse (AB_2313585) from Jackson Immuno-Research 1:10,000 for 1 hr room temperature.

### Immunofluorescence

#### Cells

IMCDs were plated on Corning 6-well 0.4 µM pore transwell inserts or Nunc-Lab Tech II Chamber Slides (Thermo) and grown to confluency. Cells were then stimulated with isotonic FBS-free control or isotonic FBS-free NaCl conditioned medium per above. After 3 hrs of stimulation, cells were washed in ice-cold PBS, fixed with 4% PFA for 30 min at RT, blocked with 5% BSA in PBS 0.2% Tween for 1 hr RT and probed with anti-aquaporin E2 FITC conjugated sc-515770 (Santa Cruz) 1:100 overnight, anti-pre-pro-vasopressin (PhosphoSolutions) 1:1000 1hr room temperature, anti-Rab3a 1:100 1hr RT (MA5-27162 – Thermo), and ActinRed555 (Life technologies) per manufacturers protocol. Cells were imaged on Nikon TiE fully motorized inverted fluorescent microscope or a Zeiss LSM 980 Confocal AiryScan 2.

#### Kidneys

Following euthanasia, kidneys were removed and incubated at room temperature overnight in 3.7% formaldehyde, 10 mM sodium m-periodate, 40 mM phosphate buffer, and 1% acetic acid. The fixed kidney was dehydrated through a graded series of ethanol, embedded in paraffin, sectioned (5 µm), and mounted on glass slides. Immunostaining was carried out as described previously (42). Primary antibodies used: anti-pre-pro-vasopressin (PhosphoSolutions) 1:1000 1hr room temperature, anti-aquaporin E2 FITC conjugated sc-515770 (Santa Cruz) 1:100 overnight, anti-Rab3a 1:100 1hr RT (MA5-27162 – Thermo), or Dolichos Biflorus Agglutinin (DBA) – Rhodamine labeled (Vector) and Lotus Tetragonolobus Lectin (LTL) – Fluorescein labeled, or ActinRed 555 ReadyProbes (Thermo). Secondary antibody used: goat-anti-Rabbit AF647 coupled (Invitrogen - A27040) and goat-anti-Mouse AF488 coupled (Invitrogen – A11001). 3D images were assembled using the Image J 3D viewer.

### Human tissue samples

All human tissue samples were obtained in accordance with and following the Vanderbilt University Medical Center IRB. Human tissue samples were obtained as unidentified frozen human tissue procured through the Collaborative Human Tissue Network (CHTN) and the formalin fixed paraffin embedded de-identified blocks procured through the biorepository of the Vanderbilt University Medical Center Department of Pathology.

### Statistical analyses

All values are expressed as mean ± SD. Between group comparisons were made using one-way ANOVA with post-hoc Tukey test or Mann-Whitney test as indicated in figure legend *p*<0.05 was used for significance threshold. Analysis was performed using Prism 9 software.

## Results

### Vasopressin mRNA is present in human and mouse kidneys

We initially interrogated the Genotype-Tissue Expression Project (GTEx) database to identify potential sites of extra-hypothalamic vasopressin production in humans (43). Vasopressin mRNA expression was highest in the hypothalamus, followed by testis, kidney cortex, and kidney medulla (Supplemental Figure 1A). We confirmed the database-defined expression of *Avp* mRNA in mouse brain and kidney tissue by performing RT-PCR on various tissues (Supplemental Figure 1B). We also found that vasopressin mRNA was present in principal cells of mice utilizing the mouse kidney single cell RNA sequencing data set available through the Kidney Interactive Transcriptomics website (http://humphreyslab.com/SingleCell) (Supplemental Figure 1C). We used the NephroSeq Application available through the University of Michigan (www.nephroseq.org) (30) to analyze publicly available, peer-reviewed, data sets which include both RNASeq and microarray data and found 22 analyses in which Avp gene expression was reported (Supplemental Table 1 and 2). To confirm these observations we created a reporter mouse, AVP-IRES2-Cre-D;ROSA^mT/mG^, in which Cre-recombinase expression linked to the vasopressin gene drives expression of membrane GFP. Cells with membrane GFP (mGFP) either actively express or are derived from cells that expressed the vasopressin gene. We found that mGFP was present in principal and intercalated cells in the AVP Cre (+) mice (Supplemental Figure 1 D – F) but not in the AVP Cre (–) mice (Supplemental Figure 1 G – I). We then analyzed single cell RNAseq data from rat primary IMCDs (44) and found that rat primary IMCDs in culture express *Avp* mRNA at levels comparable to *Aqp2*, *Aqp3*, *Aqp4*, *Scnn1a*, *Slc14a2*, and *Avpr2* (Supplemental Figure 2A,B). Together, these data suggest that the vasopressin gene is expressed in the kidney, and that collecting duct cells likely express *Avp* mRNA at baseline.

#### Hypertonicity regulates *Avp* mRNA expression in collecting duct cells *in-vitro*

As the scRNAseq data suggested there was vasopressin expression in collecting duct cells in vivo, we performed quantitative RT-PCR to explore whether *Avp* mRNA was present in mouse collecting duct cells in vitro. Vasopressin mRNA was detectable at baseline and increased when the cells were exposed to DMEM supplemented with 100 mmol of NaCl (Figure 1A, Supplemental Figure 2D). There was no statistically significant change to vasopressin mRNA expression after 12 hrs in NaCl supplemented DMEM or 24 hrs in hypotonic DMEM (Supplemental Figure 2E,F). To determine whether increased vasopressin mRNA expression was due to increased concentration of NaCl *per se* or was a response to the increased hypertonicity of the medium, we added either 100 mmol of NaCl or 200 mmol of mannitol, or glucose to the DMEM medium. The increased *Avp* transcription was only seen in the presence of NaCl supplementation, suggesting the effect was specific to the NaCl stimulation. (Figure 1B). In contrast, expression of aldose reductase (*Akr1b3*), which is sensitive to increased extracellular tonicity, increased significantly in response to both NaCl and mannitol (Figure 1C) (45). As *Aqp2* expression increases with activation of the V2r and hypertonicity (32) we measured the expression of this gene in response to NaCl, glucose or mannitol and found that only DMEM supplemented with 100 mmol of NaCl increased its expression. (Figure 1D). This increase was likely due to autocrine or paracrine vasopressin production by the IMCD cells as *Aqp2* expression was significantly inhibited by addition of the non-peptide V2r antagonist OPC-31260 (46, 47) (Figure 1E). By contrast *Akr1b3* expression induced by NaCl was not affected by the V2r antagonist (Figure 1F), suggesting that there was similar hypertonic stress between groups, and that the effect of the drug was specific. Taken together, these results indicate that cultured IMCD cells produce vasopressin mRNA, which is regulated by NaCl induced hypertonicity. In addition, they suggest that IMCD cells produce locally active vasopressin that can regulate *Aqp2* expression *in-vitro*.

#### Development of an antibody that detects locally produced pre-pro-vasopressin

We next set out to confirm that kidney collecting ducts cells produced biologically active vasopressin. To do this we needed to develop novel tools as vasopressin protein is difficult to quantify. Currently vasopressin is quantified by an ELISA assay for copeptin, which is used as a surrogate for vasopressin levels (48). Moreover, the epitopes for the commercially available antibodies target end-products of pre-pro-vasopressin processing making it impossible to distinguish site of production from peripheral tissue uptake. To confirm that kidney collecting duct cells produced biologically active vasopressin, we developed a custom antibody in which the epitope spans the known cleavage sites for the vasopressin precursor peptide (Figure 2A), which would allow us to measure locally produced pre-pro-vasopressin (49). We characterized the antibody by performing a Western blot on brain from a wild-type mouse in the presence and absence of the blocking peptide. The antibody detected an intense band at the expected weight for pre-pro-vasopressin (∼20kDa) in the brain and the signal was abolished with the blocking peptide (Supplementary Figure 3A). We then confirmed the antibody did not detect the nine amino acid variants of vasopressin or oxytocin (Supplemental Figure 3B). Immunostaining of mouse brain sections showed an intense signal in the supraoptic nucleus, containing vasopressin-producing cell bodies, and the internal layer of the median eminence, containing axons of vasopressin-producing neurons that project to the posterior pituitary (Supplemental Figure 3C) (50). To test the specificity of our antibody in cells, we transfected HEK-293 cells with a vasopressin expressing vector (pCMV6-XL-5-*AVP*) and found a single band at the expected weight for pre-pro-vasopressin (Supplemental Figure 3D). We further characterized the antibody by performing a Western blot on brain, kidney, and plasma. The antibody detected an intense band at the expected weight for pre-pro-vasopressin (∼20kDa) in the brain and the kidney (Figure 2B), but not in the plasma. To confirm the presence of pre-pro-vasopressin in the kidneys in vivo, we obtained *wild-type* mouse whole kidney lysates and incubated them with anti-pre-pro-vasopressin primary antibody with and without incubation with the blocking peptide. We detected a band of the expected weight and the signal was eliminated by the blocking peptide (Figure 2C). To verify that IMCD cells produced pre-pro-vasopressin, we performed immunoblots on cell lysates. We saw the expected single 20kDa band that decreased when the IMCDS cells were transfected with *Avp* siRNA (Figure 2D). Immunoprecipitation (IP) from whole-cell IMCD lysates using our novel antibody identified a single band at the expected weight for pre-pro-vasopressin (Supplemental Figure 3 E,F) Additionally, immunostaining of our IP sample with commercially available antibodies that target the nine-amino acid vasopressin and neurophysin-2 showed a band at the expected weight for pre-pro-vasopressin (Supplemental Figure 3G,H). Thus, we had produced a highly specific antibody that could identify un-cleaved pre-pro-vasopressin.

**Figure 2.**
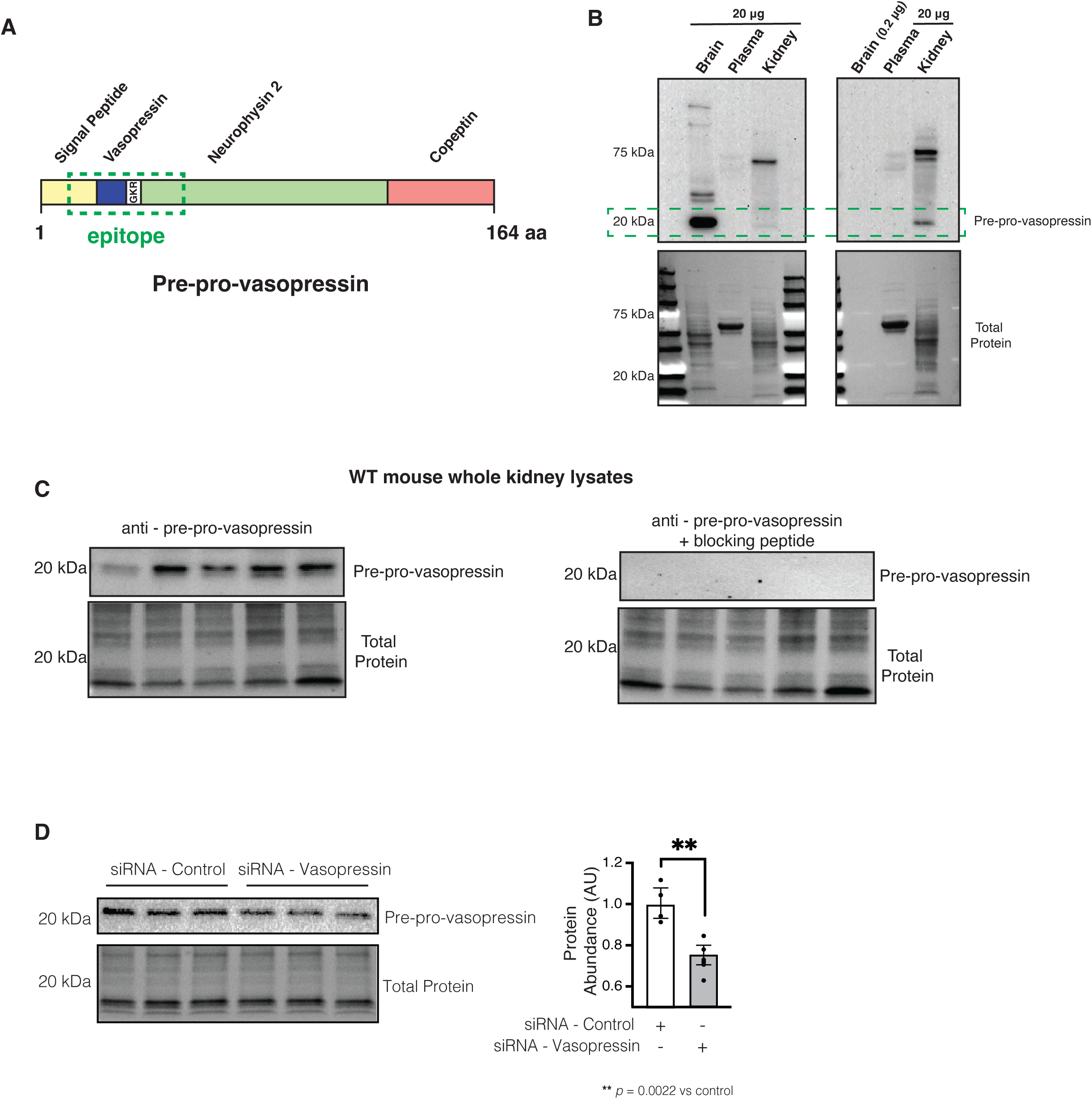
Pre-pro-vasopressin was found in mouse brain and kidney. Our novel custom antibody targets the cleavage site of the vasopressin precursor peptide (**A**). The antibody detected a specific band at 20 kDa in mouse brain and kidney samples at a 1:1 ratio (left) and a 1:10 ratio (right), but not in plasma (**B**). We detected a specific band in WT mouse kidneys (n=5), and the signal was inhibited if the primary antibody was incubated with the blocking peptide (**C**). Vasopressin was found in IMCD cells and expression was decreased with the use of siRNA that targets *Avp* (**D**). Data in D is presented as mean ± SD and analyzed with an unpaired Mann-Whitney test n=4 independent replicates ** p=0.002

#### IMCDS cells produce biologically active vasopressin

To assess if pre-pro-vasopressin protein increased in the same conditions as vasopressin mRNA, we stimulated IMCDs with NaCl for 24 hrs and determined that indeed vasopressin precursor protein expression increased (Figure 3A). Immunofluorescence staining of IMCDs stimulated with NaCl also showed increased staining for vasopressin (Figure 3G) vs controls (Figure 3D) and the signal was once again abolished with the blocking peptide (Figure 3J).

**Figure 3.**
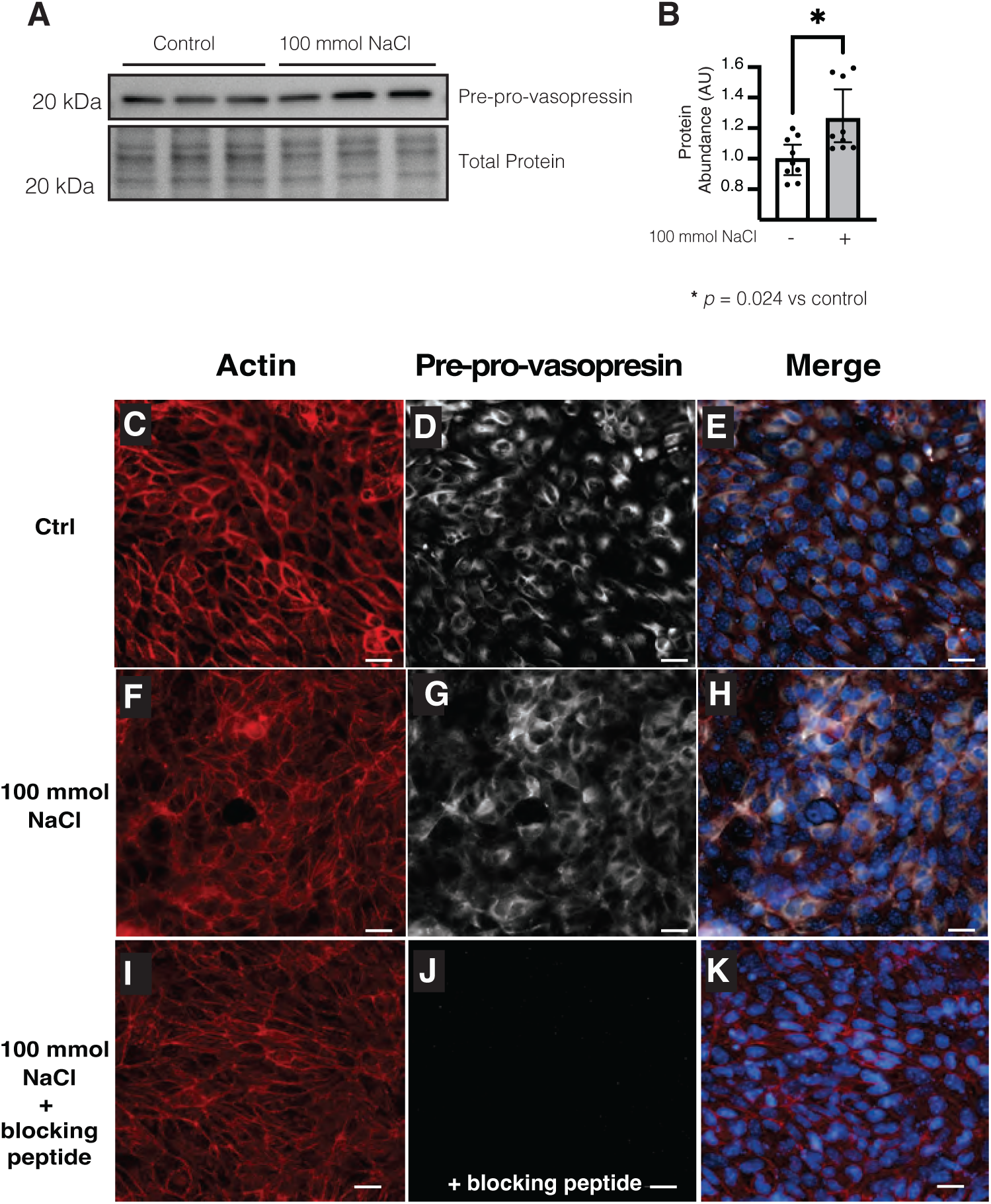
NaCl increased pre-pro-vasopressin protein in IMCD cells. Treatment of inner medullary collecting duct cells (IMCD) with NaCl for 24 hrs increased the abundance of vasopressin by immunoblot (**A,B**) and immunofluorescence (**C-K**) i. Pre-incubation of the primary antibody with the blocking peptide abolished the signal (**I-K**). Data in B are presented as mean ± SD and analyzed with an unpaired Mann-Whitney test, n=9 independent replicates * p=0.024

**Figure 4.**
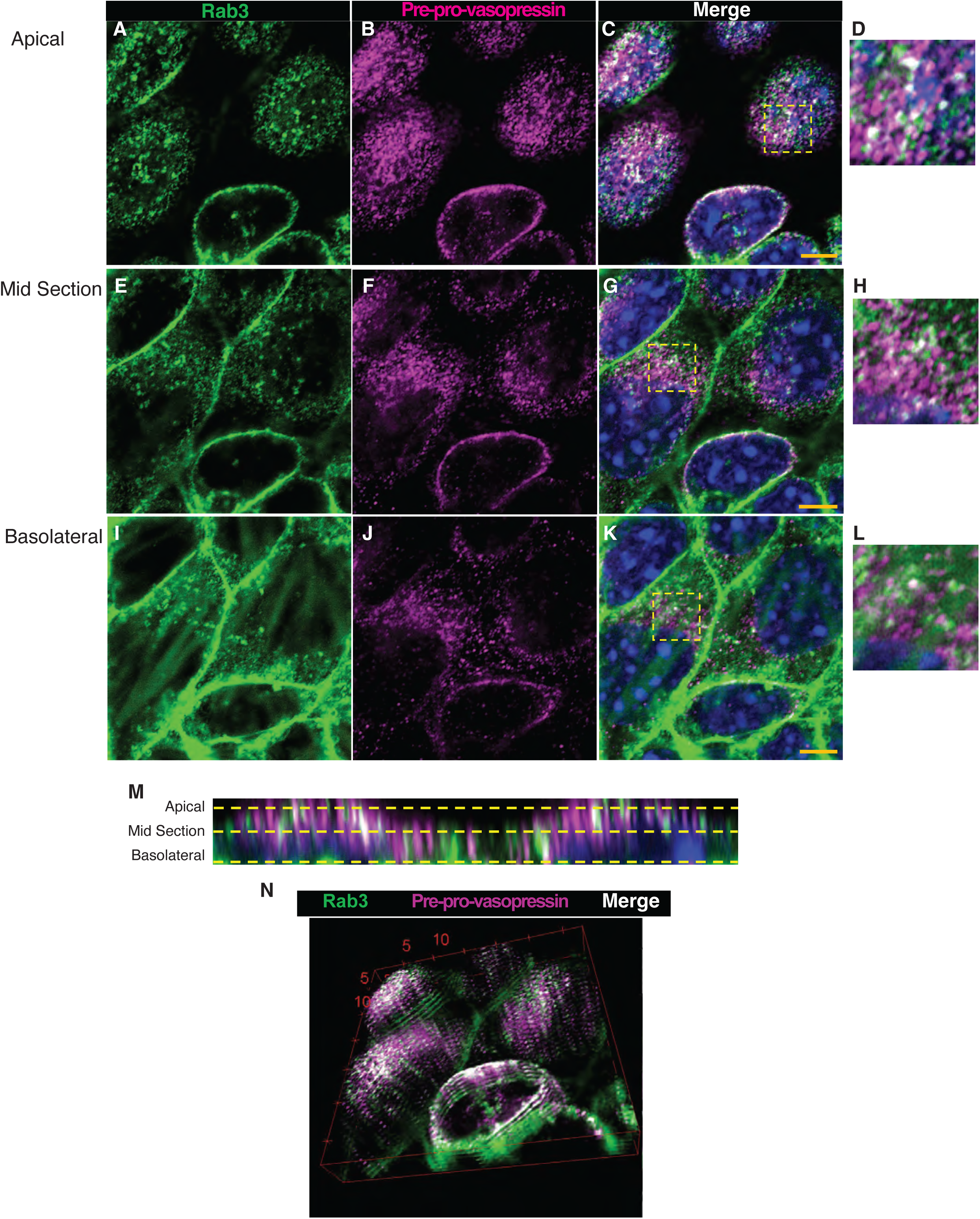
Pre-pro-vasopressin is found in Rab3 positive vesicles in vitro. Super resolution confocal images of inner medullary collecting duct cells (IMCD) stained for pre-pro-vasopressin and Rab3 show co-localization (white) of pre-pro-vasopressin (pink) in Rab3 positive vesicles (green) (**A – K**). A 3-D reconstruction of A-C shows predominantly apical co-localization of pre-pro-vasopressin and Rab3 (**M,N**).

In neurons, vasopressin is stored in secretory vesicles. Therefore, to evaluate if pre-pro-vasopressin was found in secretory vesicles in the kidney, we stained IMCDs for pre-pro-vasopressin and Rab3 and imaged with super-resolution confocal microscopy (Figure 4). We found that some, but not all pre-pro-vasopressin was colocalized with Rab3 (Figure 4 A-C) and a composite 3D image of the cells suggested apical enrichment of Rab3 positive pre-pro-vasopressin containing vesicles (Figure 4D). These data suggest that pre-pro-vasopressin in IMCDs is found in Rab3 positive secretory vesicles.

To confirm that the vasopressin produced by the IMCDS cells was biologically active we developed a novel cell line; stably transfected HEK-293 cells with the human V2 receptor (hV2R) and a cAMP response element driving luciferase expression (HEK-hV2R-CRE-Luc) that can be activated by V2 receptor activation. We then developed a bioassay in which we collected medium from control or NaCl stimulated IMCDs, illustrated in (Figure 5A). Serum starved HEK-hV2R-CRE-Luc cells were then subjected to the following: 1) no medium change, 2) fresh FBS-free DMEM, 3) fresh FBS-free DMEM + 100 mmol NaCl, 4) IMCD control medium, 5) IMCD NaCl medium, and 6) IMCD NaCl medium + OPC31260 (Figure 5B). We found that medium from IMCDs stimulated luciferase production. The luciferase activity was higher in media supplemented with NaCl than IMCD control medium, and the increased activity was abolished with the addition of OPC-31260 (Figure 5C). Together, these data suggest that there is biologically active vasopressin in IMCD conditioned medium at baseline which increases after NaCl treatment, and the NaCl dependent increase in luciferase activity was due to V2R activation as it was prevented with V2R antagonist OPC-31260.

**Figure 5.**
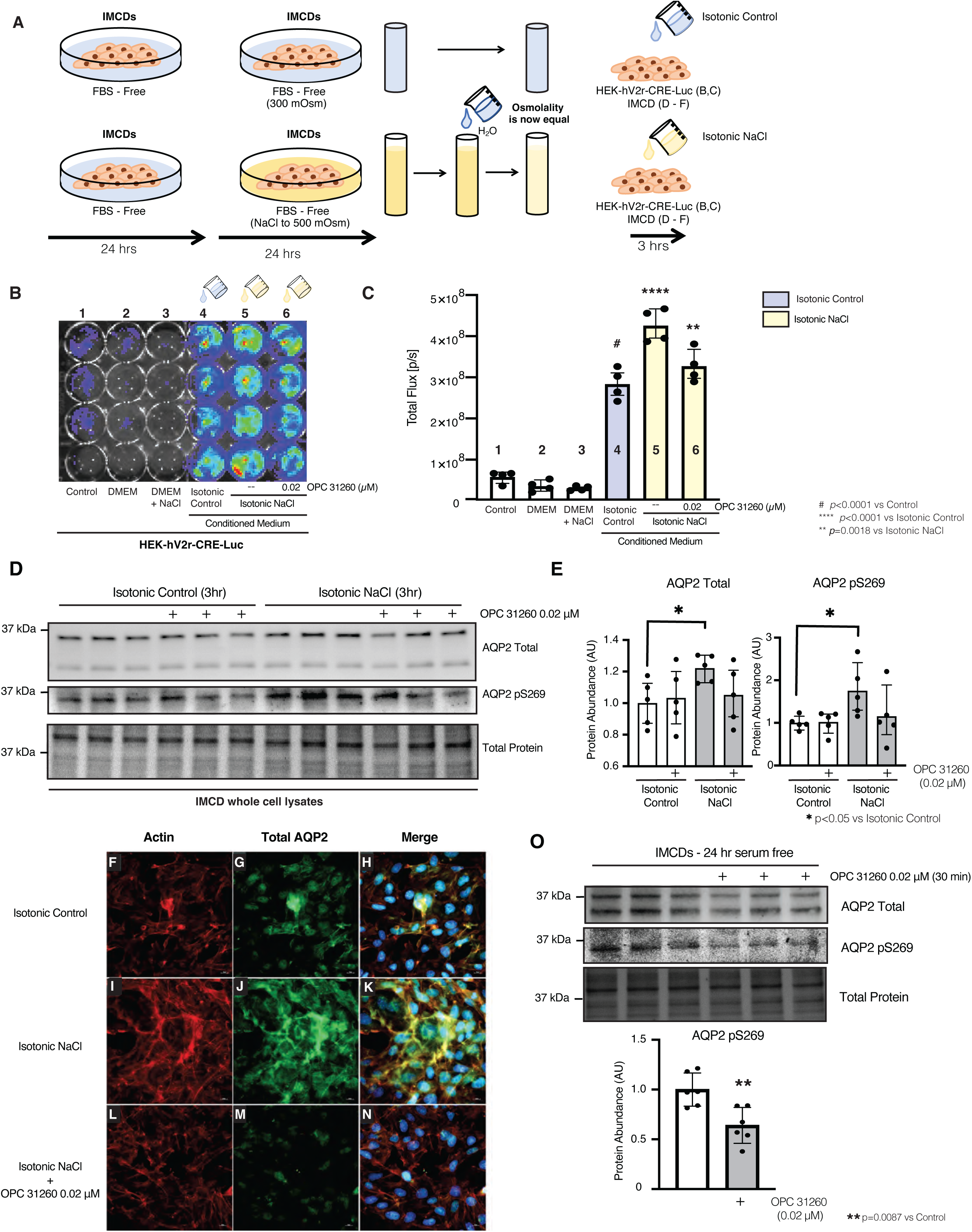
IMCD cell medium contains a V2R stimulating ligand. Bioassay experimental design to obtained conditioned medium from treated and untreated inner medullary collecting duct cells (IMCD) (**A**). HEK-V2R-Luc cells stimulated with control and NaCl treated IMCD medium had increased luciferase activity (**B,C** lanes 4,5). The NaCl mediated increase was blocked with the V2R antagonist OPC-31260 (**B,C** lane 6). Addition of isotonic + NaCl conditioned medium increased both phosphorylation of serine 269 and total abundance of AQP2, this effect is prevented with OPC-31260 (**D, E**). In IMCDs (**F-N**), isotonic + NaCl treated medium increased the staining of aquaporin 2 (AQP2) (green), relative to isotonic control medium (**G** vs **J**). The increase was prevented with OPC-31260 (**M**). Treatment of serum-starved IMCDs with OPC-31260 decreased phosphorylation of AQP2s-269 (**O**). Data in C and E are presented as mean ± SD of total flux (photons per second). Data analyzed with 1-way ANOVA with Tukey’s post hoc analysis, with a minimum of 4 independent replicates per group # p<0.0001 vs control, **** p<0.0001 vs isotonic control, ** p=0.0018 vs Isotonic + NaCl, and * p<0.05 vs Isotonic Control.

We further confirmed the presence of biologically active vasopressin in IMCD conditioned medium by adding control conditioned medium or NaCl supplemented conditioned medium (Figure 5A) to serum starved IMCDs. We then assessed the expression and phosphorylation of AQP2 (Figure 5 D-O)). As expected, when serum starved IMCDs were treated with conditioned medium from IMCDs exposed to DMEM + 100mmol of NaCl expression and phosphorylation of AQP2 increased (Figure 5 D, G, J), and this increase was blocked by the addition of OPC-31260 (Figure 5D, J, M). To directly evaluate V2r activation by locally produced vasopressin, fresh FBS-free DMEM was added to 24 serum-starved IMCDs followed by treatment with vehicle or V2r antagonist OPC-31260 for 30 minutes. Treatment with OPC-31260 decreased AQP2 phosphorylation at baseline, (Figure 5O). This suggests that baseline V2r activation in serum-starved IMCDs is dependent on local vasopressin production which can be antagonized by treatment with a V2r antagonist. Together, these data suggest that IMCD cells make biologically active vasopressin that increases when stimulated with NaCl, and local vasopressin can regulate AQP2 total protein abundance and phosphorylation.

#### Vasopressin mRNA and protein increased with water restriction *in vivo*

To confirm expression of vasopressin by the kidneys in vivo, we looked for evidence that vasopressin mRNA is made by collecting ducts in mice. We used RNA in-situ hybridization (RNAScope) and found that *Avp* mRNA colocalizes with *Aqp2* mRNA in mouse kidneys (Figure 6 A-C). This supported the publicly available RNAseq data that suggested expression of *Avp* mRNA by collecting duct cells in vivo (Supplemental Figure 2).

**Figure 6.**
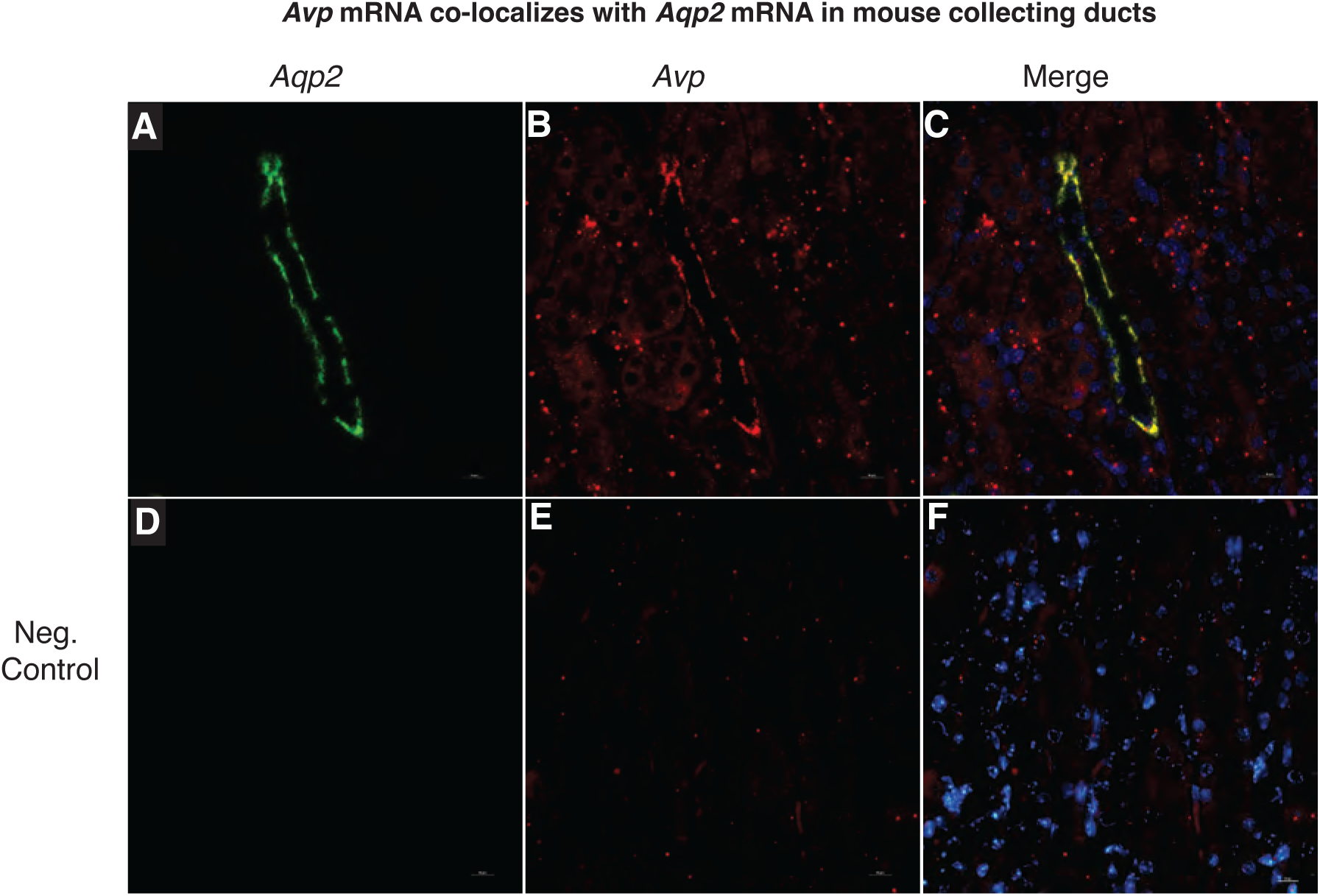
Vasopressin mRNA is found in mouse collecting duct cells in vivo. – RNA in situ hybridization (RNAScope) in mouse kidneys shows co-localization of Aqp2 (A) and vasopressin mRNA (B) in collecting ducts (C). Negative control (D-F). Scale bar = 5 uM.

To test whether expression of vasopressin mRNA in the kidney is regulated in vivo, we water restricted, and water loaded *wild type* mice for 24 hrs. We then assessed whole kidney *Avp* mRNA by qPCR and ddPCR, protein for pre-pro-vasopressin expression, and confirmed localization via immunofluorescence. Both vasopressin mRNA and protein were higher in water restricted vs water loaded mice (Figure 7 A-C, Supplemental Figure 4). Consistent with our in vitro data, pre-pro-vasopressin protein was found in collecting ducts (Figure 7 D-K). Moreover, collecting duct pre-pro-vasopressin signal was higher in water restricted vs water loaded mice (Figure 7 F, J).

**Figure 7.**
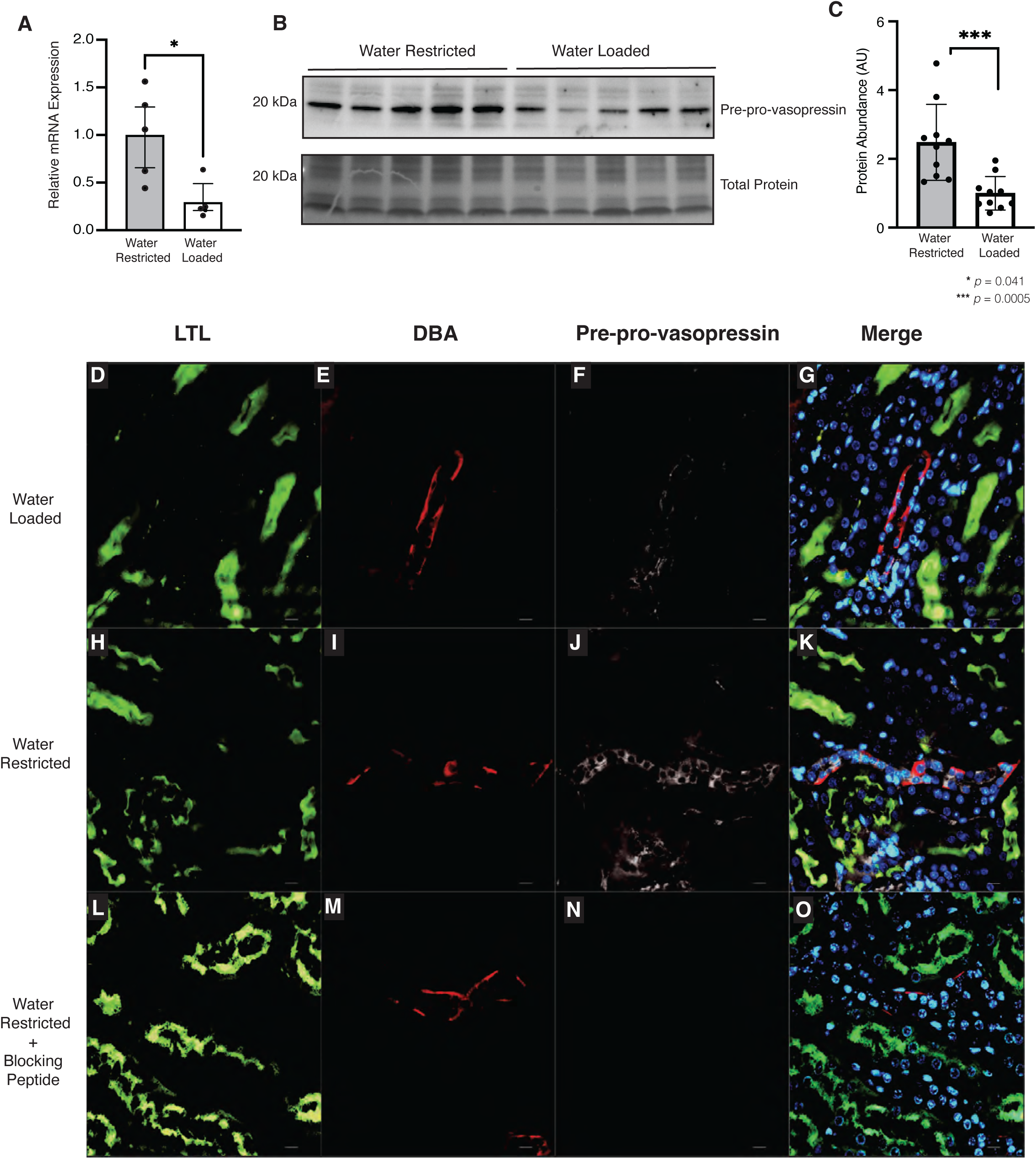
Kidney-derived vasopressin increases after water restriction. *Avp* mRNA increased in water restricted mice relative to water loaded controls (**A**). Kidney-derived pre-ppro-vasopressin protein increased after water restriction relative to water loading (**B**). In mouse kidney (**D-O**) Kidney-derived pre-pro-vasopressin signal (white) in collecting ducts (DBA – red) increased in water restricted animals (**J vs F**) and the signal was abolished with primary antibody pre-incubation with blocking peptide (**N vs J**). Data in A and C are presented as mean ± SD and analyzed with an unpaired Mann-Whitney test, * p=0.041, ** p = 0.013, water restricted n=10, water loaded n=10.

#### Vasopressin protein is made by human kidneys

Whole kidney mRNA data (Supplemental Figure 1A) suggested vasopressin expression in human kidneys. Therefore, we obtained samples of frozen, and formalin fixed paraffin embedded human kidney tissue to test for vasopressin protein expression. We found that human kidneys can express vasopressin mRNA (Supplemental Figure 4B) which co-localizes with *AQP2* mRNA (Supplemental Figure 5) and pre-pro-vasopressin protein (Figure 8A). Moreover, pre-pro-vasopressin is found in collecting ducts (Figure 8 B-F) and it co-localizes with Rab3 in the apical surface of CD cells (Figure 9). Together, these data indicate that vasopressin is found in human CD cells in vivo.

**Figure 8.**
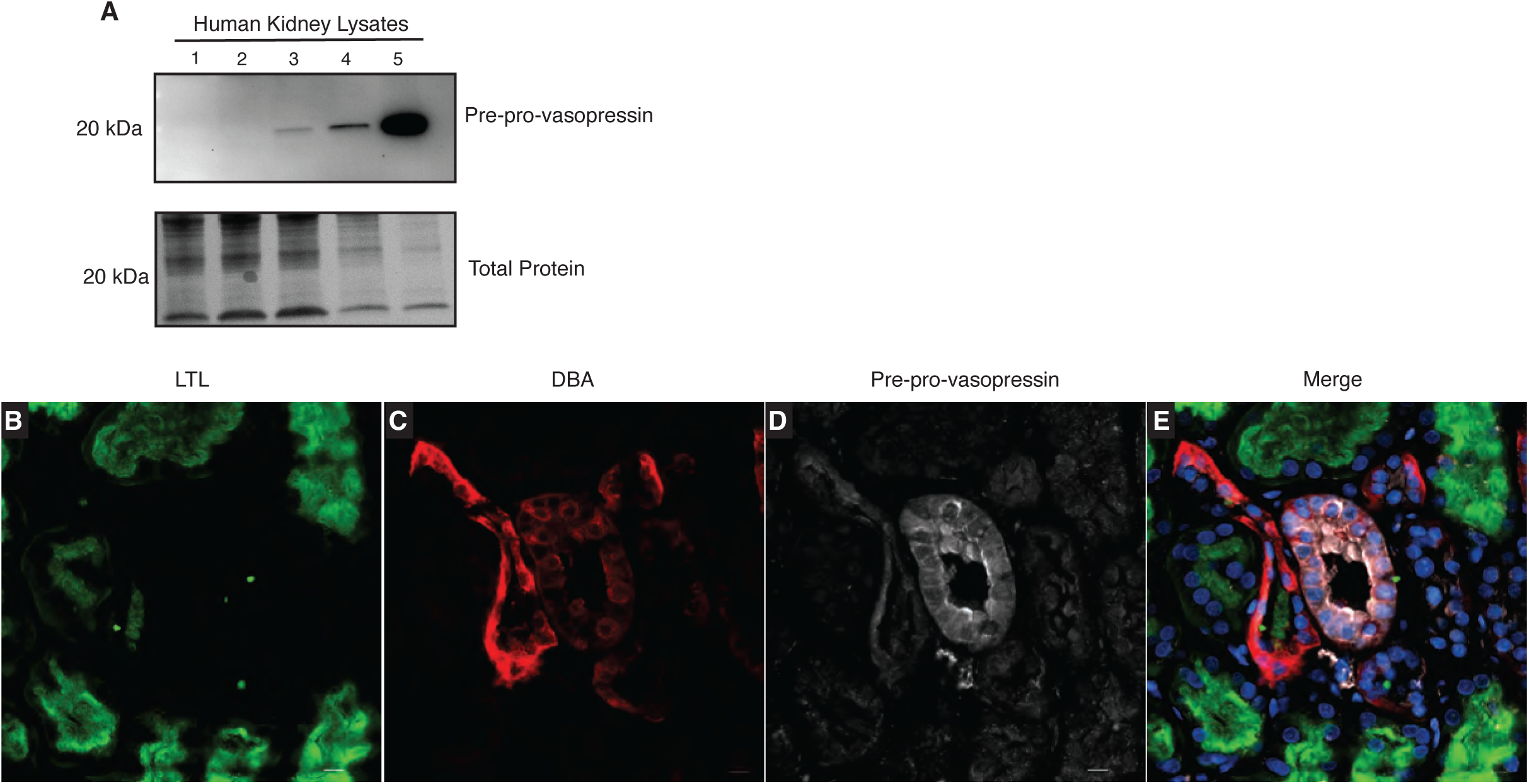
Human kidneys make vasopressin. Pre-pro-vasopressin was found in healthy human kidney lysates (A). Pre-pro-vasopressin (white) was found in the collecting ducts (DBA – red), but not in the proximal tubules (LTL – green) (**B – E**). Scale bar =10 uM.

**Figure 9.**
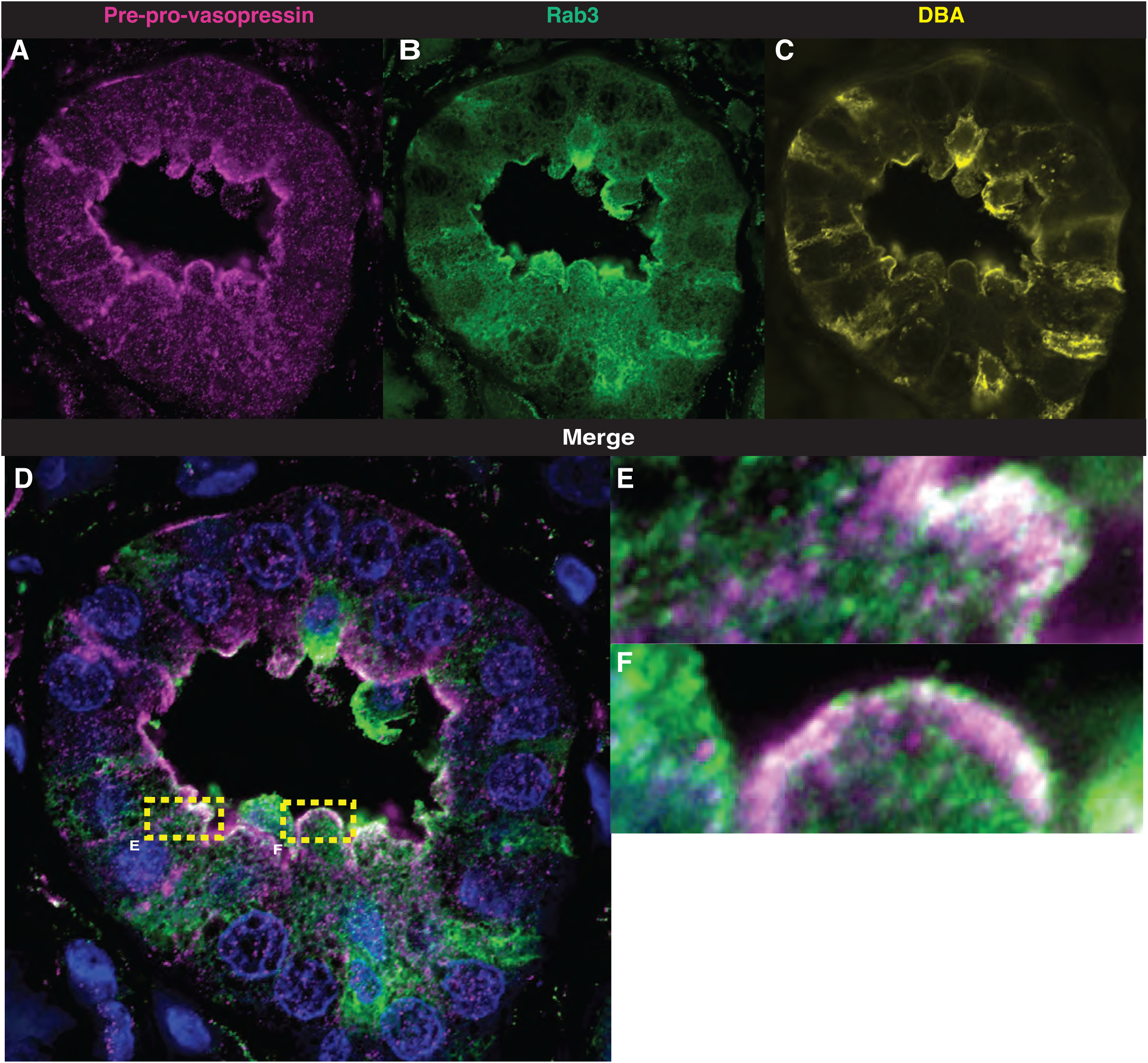
Pre-pro-vasopressin co-localizes with Rab3 in human collecting ducts. Super-resolution confocal imaging shows that pre-pro-vasopressin (pink) co-localizes with Rab3 (green) in DBA (yellow) stained human collecting ducts (**A-F**). Co-localization (white). Field in A-D is 74 uM x 74 uM.

## Discussion

Vasopressin is thought to be made primarily in the brain, and that this is the sole source of vasopressin stimulating vasopressin V2 receptors in the kidney. In the present study we challenge the current dogma. We developed a novel antibody that recognizes the pre-pro-vasopressin peptide to study local production of vasopressin in the kidney. We used this antibody to confirm that pre-pro-vasopressin was made in mouse collecting ducts cells in vitro and in vivo. We then showed that kidney-derived pre-pro-vasopressin production was responsive to water restriction, which increased mRNA and protein expression in mice. Finally, we also determined that vasopressin mRNA and pre-pro-vasopressin protein were found in healthy human kidney collecting ducts. Thus, we provide novel evidence of physiologic biologically active extra-hypothalamic production and regulation of kidney-derived vasopressin from kidney epithelial cells.

Given the prior reports of vasopressin production outside of the brain (21-29), and the publicly available scRNA-sequencing data, we confirmed vasopressin mRNA in mouse and human kidneys (Supplementary Figure 1, Supplementary Figure 2, Figure 6, Supplementary Figure 5) in collecting duct cells. Our reporter mouse further confirmed the expression of *Avp* mRNA in vivo in collecting duct epithelial cells. Although the presence of mGFP could mean *Avp* gene expression occurred in a progenitor cell, our data showing changes in *Avp* mRNA abundance in vivo suggest that there is ongoing *Avp* gene expression in kidney collecting duct cells.

In the hypothalamus, *Avp* can be stimulated by NaCl (51). We found that in collecting duct cells in vitro, *Avp* is expressed at baseline in vitro and that NaCl increases *Avp* mRNA (Figure 1). Interestingly, NaCl is known to increase *Aqp2* independent of exogenous vasopressin, but the contribution of endogenous vasopressin had never been studied (5, 32). We found that V2r was activated in vitro by endogenous vasopressin in response to hypertonic stress. Moreover, the addition of a V2r antagonist prevented hypertonicity induced increase in *Aqp2* expression in a dose dependent manner (Figure 1E). Thus, these data suggest that vasopressin is produced by collecting duct cells in vitro and plays a role regulating *Aqp2* expression.

Vasopressin is difficult to quantify (52, 53). Commercially available methods to measure vasopressin including ELISAs and antibodies all detect terminally processed peptides, preventing discrimination between local production versus peripheral uptake of hypothalamic vasopressin or related peptides. To circumvent this, we developed an antibody that specifically detects the pre-pro-vasopressin peptide, as the epitope spans the cleavage regions (Figure 2A). Our antibody detected a band at the expected weight (∼20 kDa) in mouse brain and kidney (Figure 2B, Supplemental Figure 3A), it stained regions of the brain known to produce vasopressin (Supplemental Figure 3C) as well as murine IMCDs (Figure 3). Moreover, after IP with our antibody, separate anti-vasopressin and anti-neurophysin 2 antibodies detected a band at the expected weight for pre-pro-vasopressin (Supplemental Figure 3 E-H). With this novel antibody, we were able to determine that immunoreactive pre-pro-vasopressin can be found in both mouse and human whole kidney lysates (Figure 2C,D Figure 8A,B), localizes to the distal nephron, and can be found in Rab3 positive vesicles in mice and humans, which suggests it can be secreted (Figures 4 and 9). To our knowledge this is the only available method that can detect the vasopressin pre-pro-peptide. This is critical, as peripheral uptake of circulating vasopressin or copeptin could account for non-specific staining with other methodologies.

We designed a bioassay to assess if the locally produced pre-pro-vasopressin was biologically active (Figure 5A) and found that biologically active vasopressin was present in medium from IMCDs (Figure 5B-O). Our bioassay relied on V2r activation by a ligand found in IMCD supernatant to assess the biologic activity of vasopressin. In our model, it possible that luciferase and/or V2R being activated independent of vasopressin. However, the decrease in luciferase signal (Figure 5B lanes 5-6) and AQP2 phosphorylation (Figure 5O) seen with the addition of OPC-31260 suggests that there is a V2R specific ligand in IMCD medium. Our data does not preclude the possibility that local vasopressin could be signaling through other pathways including intracellular binding to the V2R (54). Although cancer cell lines have been reported to make vasopressin (55), and there are reports of vasopressin mRNA expression in tissues outside of the hypothalamus (21-29), the production of biologically active vasopressin by non-malignant cells has not been reported.

Hypertonicity is a known regulator of hypothalamic vasopressin production. We found that both *Avp* mRNA (Figure 7A, Supplemental Figure 4) and pre-pro-vasopressin protein (Figure 7B,C) were higher in kidneys from water restricted vs water loaded animals. These in vivo observations corroborated our in vitro data and suggest that local vasopressin could be playing a role in urine concentration and dilution in vivo. From these results we can conclude that kidney-derived vasopressin is produced by collecting duct cells in vivo and production is regulated by hypertonic stress.

The physiologic pathways through which hypothalamic vasopressin is involved in regulation of water and blood pressure homeostasis have been known for over 50 years (1). However, there is a large body of literature on the non-water non-blood pressure effects of vasopressin for which physiologic feedback loops have not been clearly defined (7-20). Non-osmotic vasopressin production has been ascribed to hypothalamic stimulation via other mechanisms e.g., nausea and pain (6, 56), but to our knowledge there is no prior reported data on vasopressin production outside of the brain under physiologic conditions. Interestingly, the vasopressin peptide evolved prior to the development of the vertebrate neurohypophyseal system (57). This finding could explain why cells outside of the neurohypophyseal system express vasopressin. As such, non-malignancy associated extra-hypothalamic production of vasopressin has also been reported previously in the heart, ovaries, testis, and adrenal glands (21-29). However, these studies did not establish whether this extra-hypothalamic vasopressin was biologically active.

Our study extends the current model in which vasopressin is solely produced in the brain under physiologic conditions. Many questions remain regarding the in vivo relevance of our observations. Given that kidney-derived vasopressin is stimulated by hypertonicity and patients with diabetes insipidus (DI) have low medullary tonicity, we would expect patients with central or nephrogenic DI to have low to absent levels of kidney-derived vasopressin. This might explain why kidney-derived vasopressin is unable to overcome the concentrating defect in patients with nephrogenic DI. Interestingly, V2R stimulation has differing effects depending on whether vasopressin is apical or basolateral, and reports of intracellular activation of the V2R open the door to the possibility that local vasopressin production may regulate V2R from within the cell (54, 58). Moreover, there are data that suggest that kidney-derived vasopressin mRNA is upregulated in humans with CKD (30, 59) (Supplemental Figure 6A) and in mice after kidney injury (60) (Supplemental Figure 6B), implying that the local kidney-vasopressin system could be involved in pathways beyond water homeostasis.

In conclusion, our data show that there is regulated expression of the vasopressin gene in the kidney in both mice and humans and kidney epithelial cells make biologically active vasopressin in response to NaCl mediated hypertonicity. It is well known that vasopressin contributes to progression of non-diabetic, diabetic, and polycystic kidney disease (18, 61-63). Our identification of a local vasopressin system in the kidney could provide insight as to how the vasopressin system contributes to kidney function in health and disease.

## Acknowledgements

The Genotype-Tissue Expression (GTEx) Project was supported by the Common Fund of the Office of the Director of the National Institutes of Health, and by NCI, NHGRI, NHLBI, NIDA, NIMH, and NINDS. The data used for the analyses described in this manuscript were obtained from the GTEx Portal on 01/15/2022. Biospecimens were provided by the NCI funded Cooperative Human Tissue Network (CHTN). Translational Pathology Shared Resource at VUMC is supported by NCI/NIH Cancer Center Support Grant P30CA068485. Graphical abstract was created with BioRender.com We are thankful for the support provided by Dr. Timothy Blackwell, Taylor Sherrill, and David Nichols.

## Funding

These studies were supported by NIH grants, DK51265, DK95785, DK62794, DK7569, P30DK114809 (RCH, MZZ), DK116964 (GB), DK069921, DK127589 (RZ) VA Merit Award 00507969 (RCH), VA Merit I01-BX002196 (RZ), ASN-Kidney Cure career development award (JAW), Intramural Research Program of the NIH, NIEHS ES103361-01 (JAW), the Vanderbilt Center for Kidney Disease, and CTSA award No. UL1 TR002243 from the National Center for Advancing Translational Sciences. Its contents are solely the responsibility of the authors and do not necessarily represent official views of the National Center for Advancing Translational Sciences or the National Institutes of Health. We acknowledge the Translational Pathology Shared Resource supported by NCINIH Cancer Center Support Grant 5P30 CA68485-19.

Juan Pablo Arroyo is a Robert Wood Johnson Foundation Harold Amos Medical Faculty Development Program Scholar Andrew Terker was supported by a postdoctoral fellowship from the American Heart Association.

Fabian Bock is supported by a postdoctoral Ben J. Lipps fellowship from the American Society of Nephrology

## Author Contributions

JPA, GB, MW, and RCH, conceived the study and designed experiments. JPA, YZ, CM, FB, MJ, RV, MK, WL, JC performed experiments. JPA, AST, FB, JAW, ERG, AXM, RV, MK, RZ, EH, MHW, GB, MZ, MW, and RCH analyzed data. JPA, AST, EH, RZ, GB, and RCH wrote and edited the manuscript.

## Conflict of Interest

The authors have declared that no conflict of interest exists

---

**Supplemental Figure 1.**
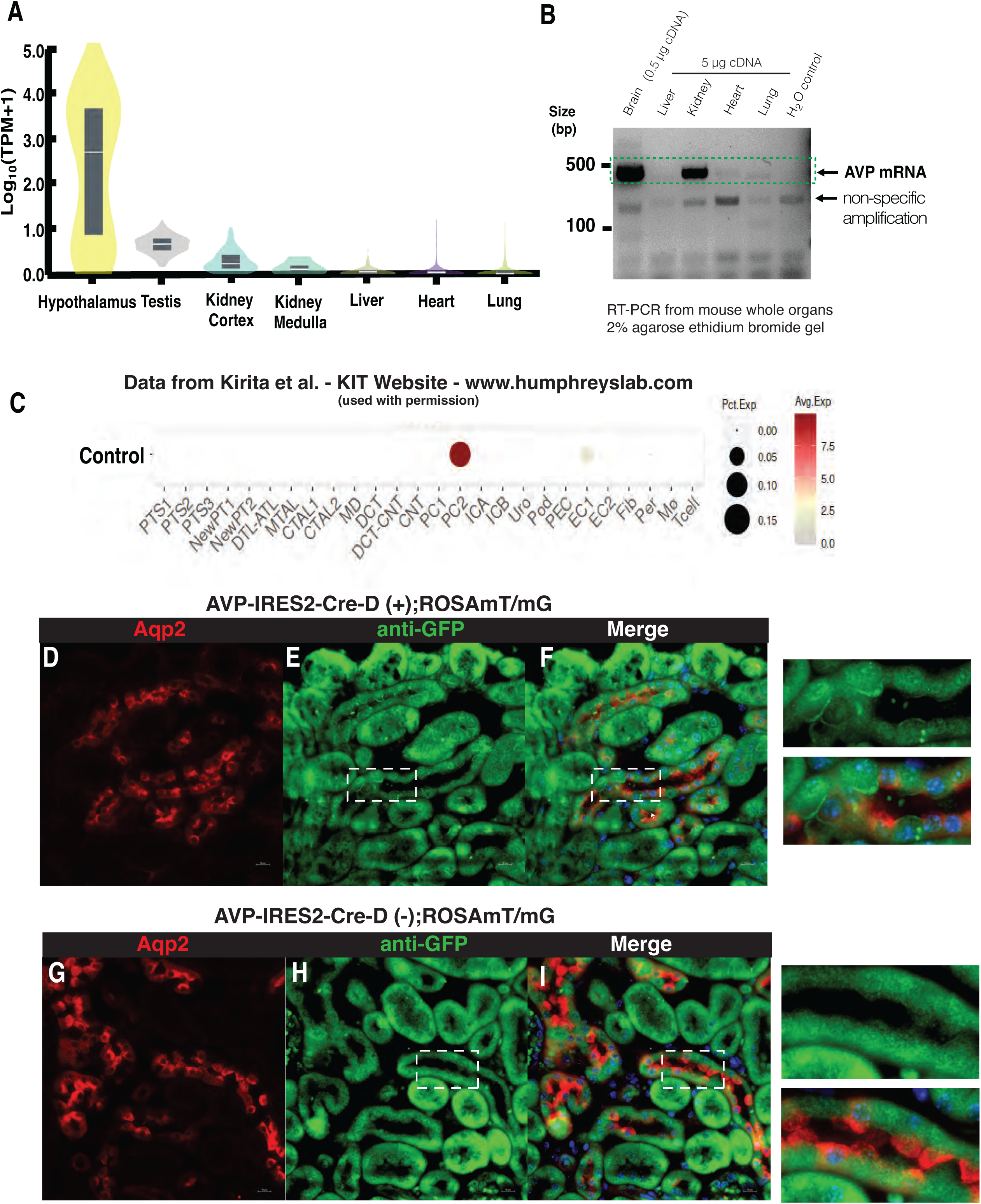
Mouse and human kidneys have vasopressin mRNA. Vasopressin gene mRNA is present in humans and mice and localizes to medullary collecting duct cells, **A)** GTEx database – bulk tissue gene expression of *Avp* in human tissue, **B)** mouse RT-PCR which shows vasopressin mRNA in brain and kidney tissue (expected size 494 base pairs), **C)** mouse kidney single cell RNAseq from the Kidney Interactive Transcriptomics (http://humphreyslab.com/SingleCell/) that shows expression of the vasopressin gene in medullary collecting duct principal cells (PC2) at baseline. (Used with permission). AVP-IRES-Cre-D(+); mTmG **(D-F)** express membrane GFP (arrows), while AVP-IRES-Cre-D(-); mTmG **(G-I)** do not. (PTS1, proximal tubule segment 1; PTS2, proximal tubule segment 2; PTS3, proximal tubule segment 3; NewPT1/PT2, proliferating proximal tubule; DTL-ATL, thin descending and ascending limb of the loop of Henle; MTAL, medullary thick ascending of the loop of Henle; CTAL1-2, cortical thick ascending limb of the loop of Henle; MD, macula Densa; DCT, distal convoluted tubule; PC1, cortical principle cell; PC2, medullary collecting duct principal cells; ICA/ICB, alpha and beta intercalated cells; Uro, urothelium; Pod, podocytes; PEC, parietal epithelial cell; EC1/2 endothelial cells; Fib, fibroblast; Per, pericytes; Mì, macrophage; Tcell, T-lymphocytes)

**Supplemental Figure 2.**
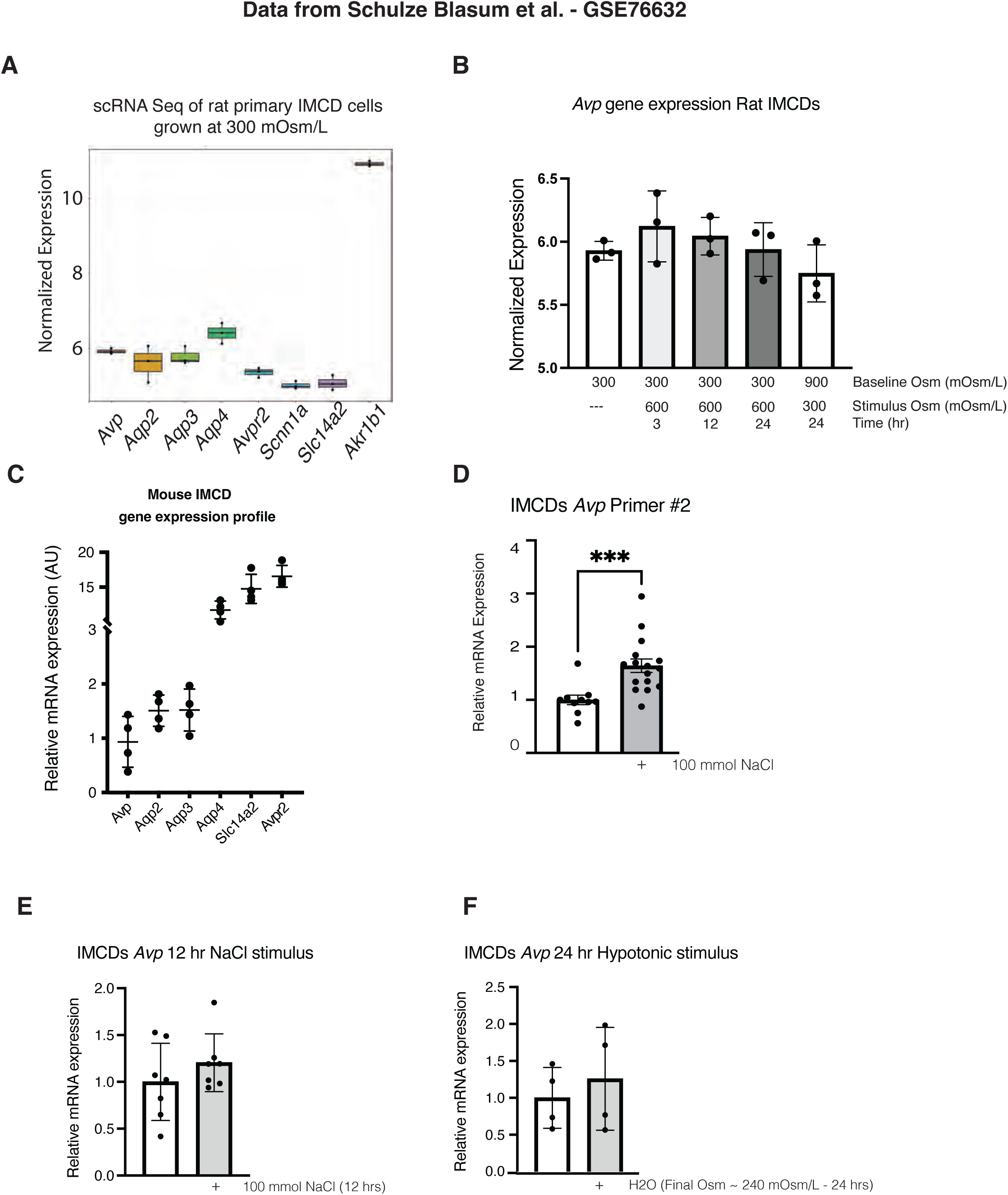
Expression of vasopressin by cells and characterization of collecting duct cell line and vasopressin mRNA expression. **A**) Single cell RNA sequencing of rat primary inner medullary collecting duct cells shows similar levels of expression of vasopressin (Avp), Aquaporin 2 (Aqp2), Aquaporin 3 (Aqp3), Aquaporin 4 (Aqp4), Vasopressin receptor type 2 (Avpr2), Epithelial Na+ Channel (Scnn1a), and Urea Transporter 2 (Slc14a2). **B**) Hypertonic stimulation causes a non-statistically significant increase in Avp mRNA expression in rat 1ry IMCDs. Our mouse inner medullary collecting duct cells (IMCDs) express the genes characteristic for IMCDs including Aquaporin 2 (Aqp2), Aquaporin 3 (Aqp3), Aquaporin 4 (Aqp4), Vasopressin receptor type 2 (Avpr2), and Urea Transporter 2 (Slc14a2) (**C**). A second primer that spans exon-exon segments also found vasopressin mRNA in IMCDs and vasopressin mRNA expression increased after stimulation with NaCl (**D**). 12 hr stimulation with the addition of 100 mmol NaCl to serum-free DMEM or 24 hr stimulation with the addition of ddH2O to serum-free DMEM (Final Osm ∼ 240 mOsm/L) for 24 hrs had no statistically significant differences in vasopressin mRNA expression **(E,F)**. Data are presented as mean ± SD and analyzed with 1-way ANOVA with Tukey’s post hoc analysis, with a minimum of 4 independent replicates per group *** p<0.001 vs control.

**Supplemental Figure 3.**
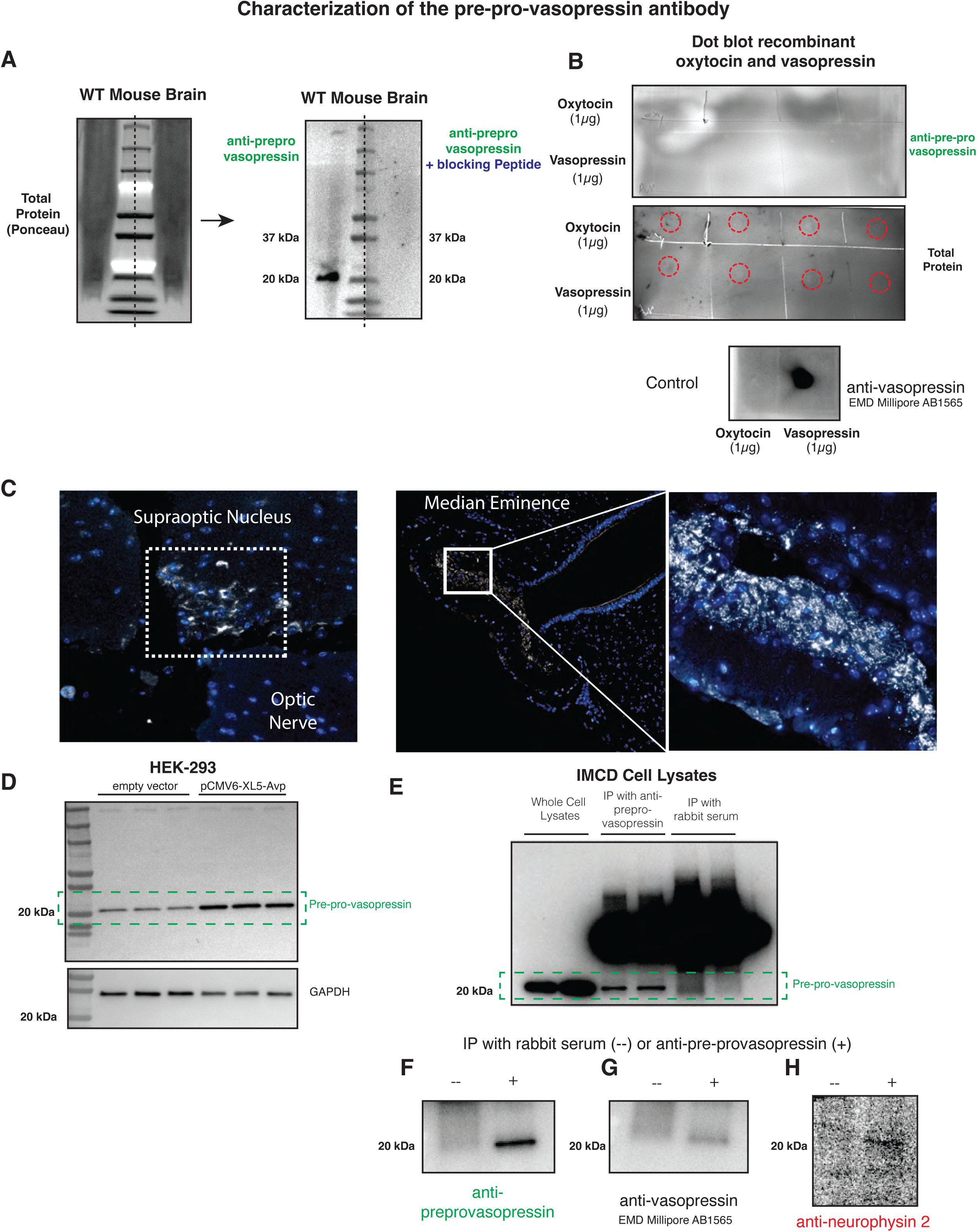
– **Characterization of our pre-pro-vasopressin antibody** Mouse whole brain lysates were run and incubated with anti-pre-pro-vasopressin alone or pre-incubated with the blocking peptide. Primary antibody alone recognized a single band at the expected weight (20kDa), and the signal was abolished when the primary antibody was pre-incubated with the blocking peptide (**A**). To corroborate that our antibody only detected the pre-pro-vasopressin we blotted 1 ug of recombinant vasopressin or oxytocin on nitrocellulose and then probed with anti-pre-pro-vasopressin primary. No signal was detected, confirming that our antibody only detects the un-processed vasopressin (**B**). We then confirmed that our antibody detected vasopressin in the expected anatomic regions in a WT mouse brain (**C**). Transfection of vasopressin expressing vector pCMV6-XL5-Avp in HEK-293 cells shows increased signal at the expected weight for pre-pro-vasopressin when probed with our novel anti-pre-pro-vasopressin antibody (**D**). Immunoprecipitation of pre-pro-vasopressin with our anti-pre-pro-vasopressin antibody or rabbit serum (negative control) from IMCD whole cell lysates (**E**) showed a band at the specific weight for pre-pro-vasopressin when probed with our antibody (**F**), an antibody against the nine amino acid vasopressin peptide (**G**), and an antibody against neurophysin 2 (**H**).

**Supplemental Figure 4.**
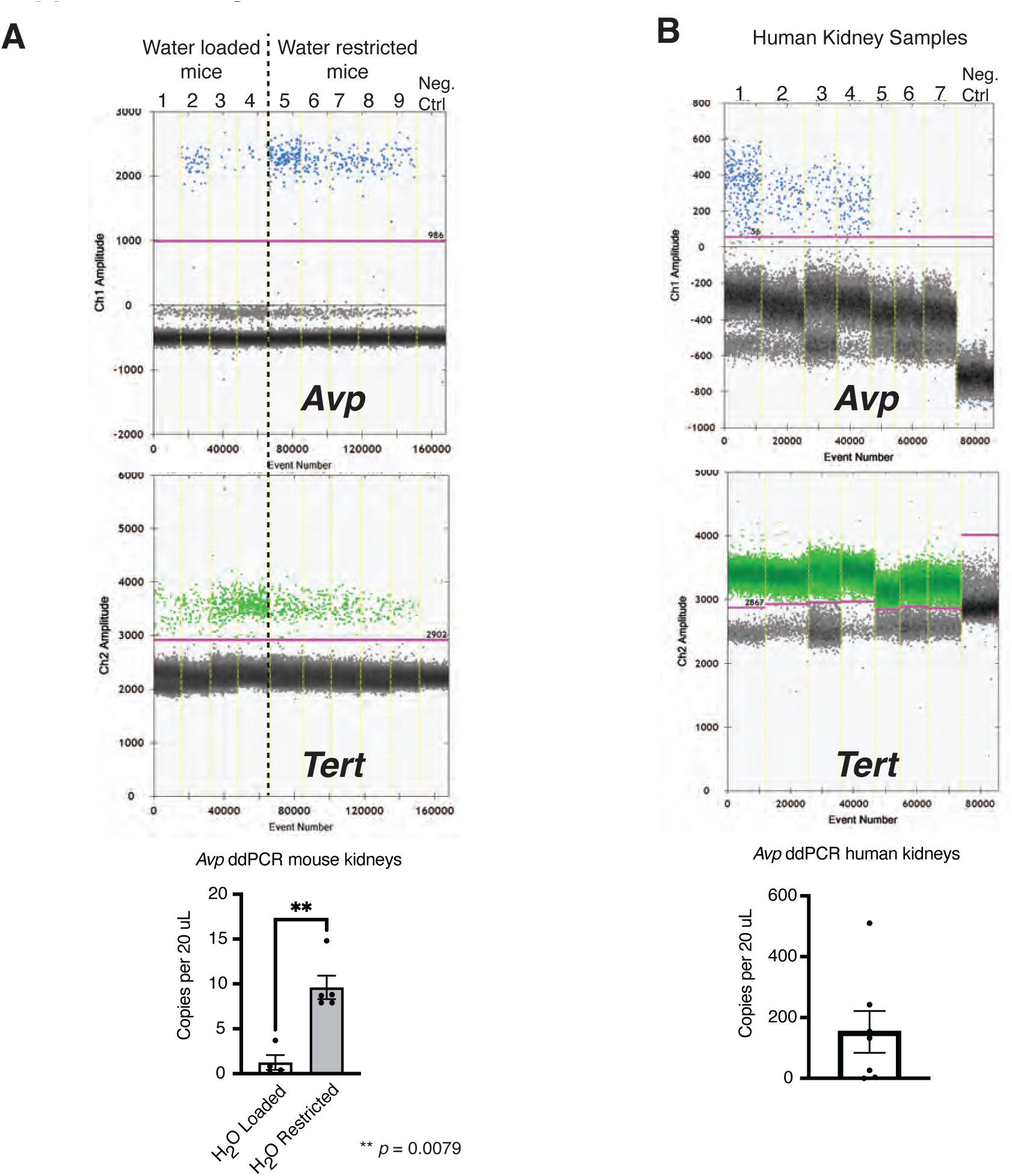
– **Droplet digital PCR (ddPCR) of mouse and human kidneys shows expression of vasopressin mRNA** Droplet digital PCR (ddPCR) of water loaded and water restricted mice (A) and human kidneys (B), shows expression of vasopressin mRNA in copies per 20 uL normalized to Tert gene expression. Data are presented as mean ± SD and analyzed with an unpaired Mann-Whitney, with a minimum of 4 independent replicates per group *** p<0.0079 vs water loaded.

**Supplemental Figure 5.**
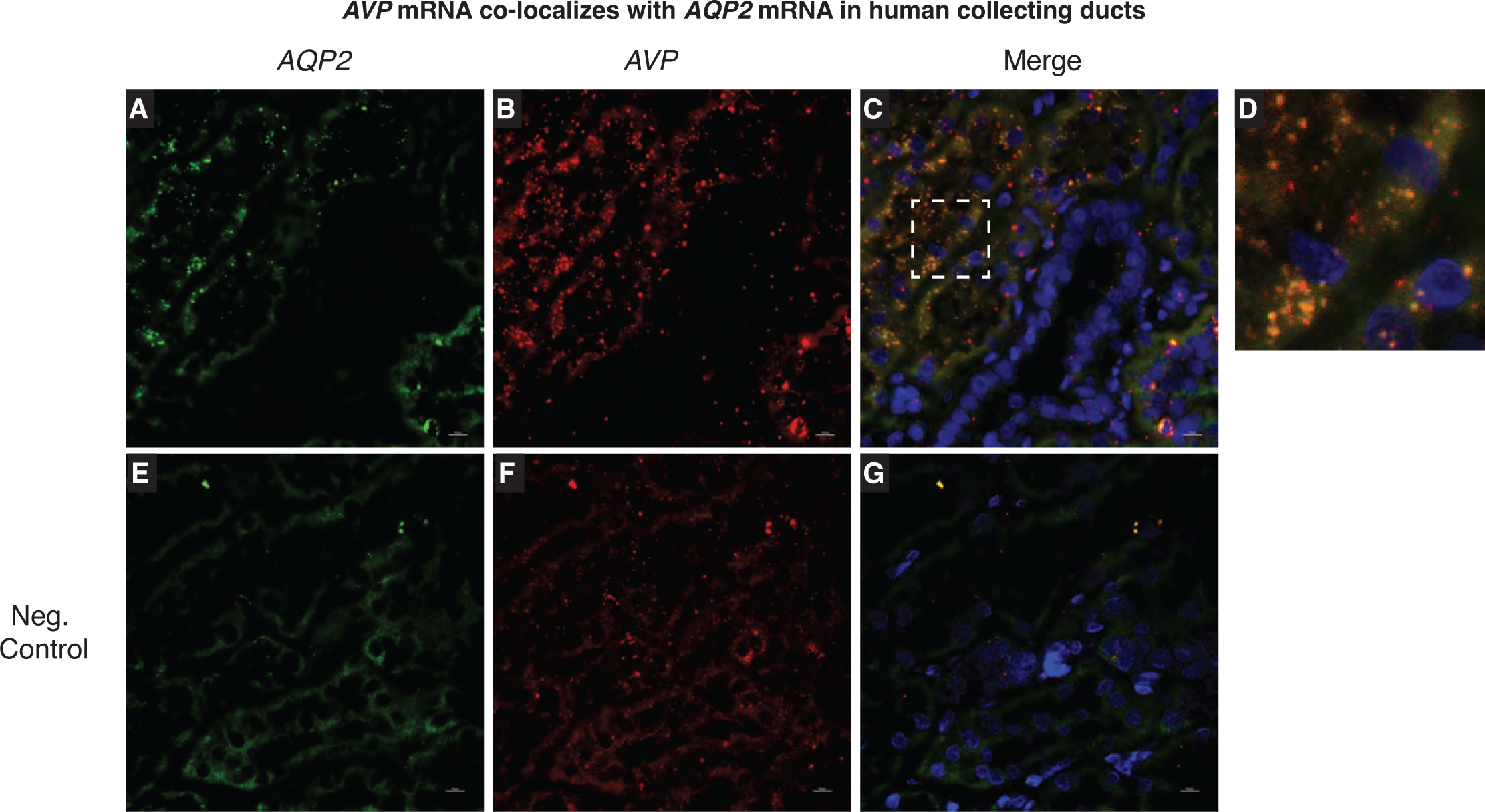
Vasopressin mRNA is found in human collecting duct cells in vivo. – RNA in situ hybridization (RNAScope) in human kidneys shows co-localization of Aqp2 (A) and vasopressin mRNA (B) in collecting ducts (C,D). Negative control (E-H). Scale bar = 10 uM.

**Supplemental Figure 6.**
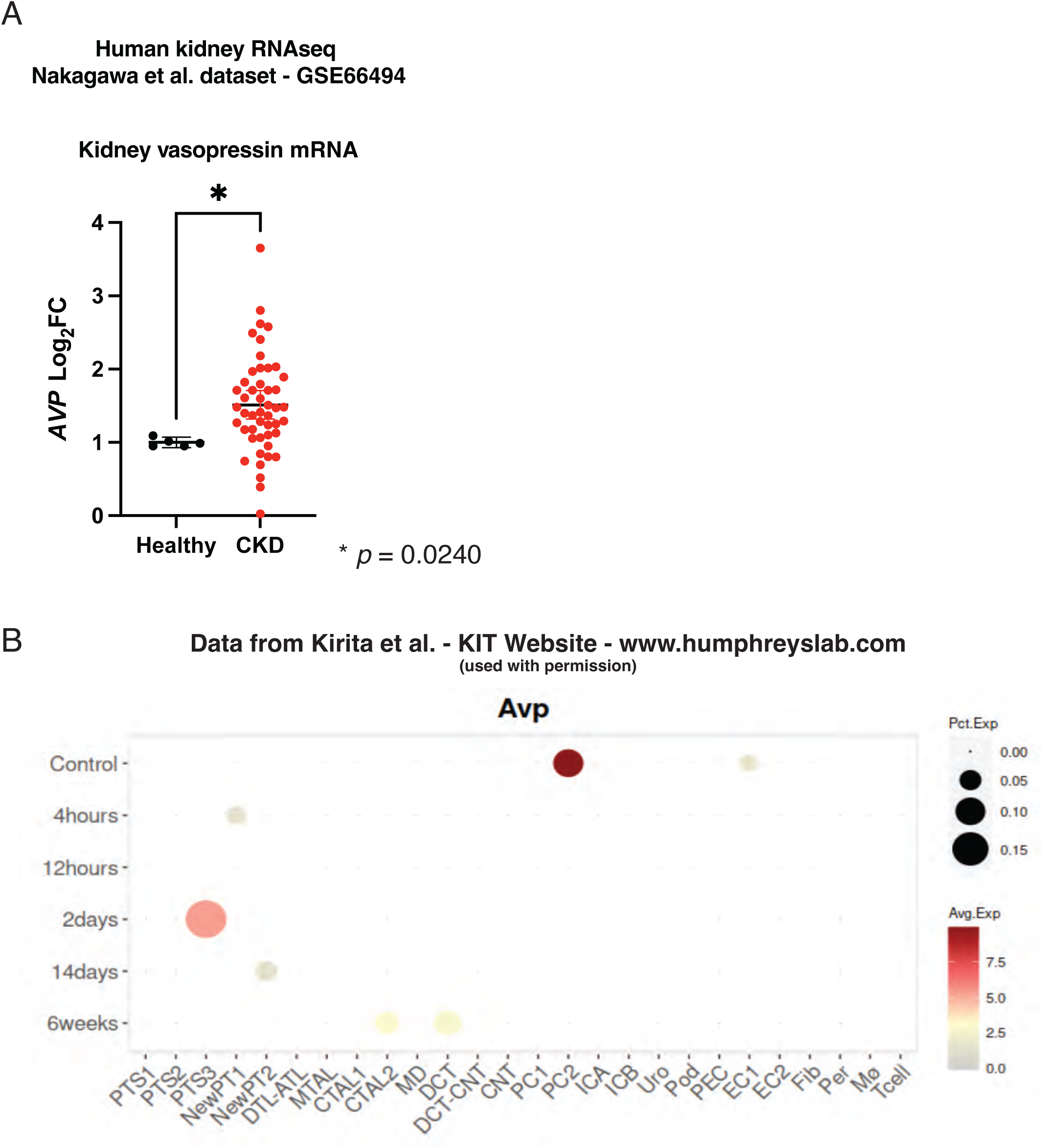
Whole kidney mRNA from non-CKD and CKD kidney biopsies shows a 1.8 fold increased expression of *Avp* mRNA in patients with CKD (**A**). Mouse kidney single cell RNAseq from the Kidney Interactive Transcriptomics (http://humphreyslab.com/SingleCell/) that shows expression of the vasopressin gene is induced in multiple nephron segments after ischemia reperfusion injury (**B**). (PTS1, proximal tubule segment 1; PTS2, proximal tubule segment 2; PTS3, proximal tubule segment 3; NewPT1/PT2, proliferating proximal tubule; DTL-ATL, thin descending and ascending limb of the loop of Henle; MTAL, medullary thick ascending of the loop of Henle; CTAL1-2, cortical thick ascending limb of the loop of Henle; MD, macula Densa; DCT, distal convoluted tubule; PC1, cortical principle cell; PC2, medullary collecting duct principal cells; ICA/ICB, alpha and beta intercalated cells; Uro, urothelium; Pod, podocytes; PEC, parietal epithelial cell; EC1/2 endothelial cells; Fib, fibroblast; Per, pericytes; Mì, macrophage; Tcell, T-lymphocytes)

**Supplementary Table 1.**

| Summary of Analyses for AVP |  |  |
| --- | --- | --- |
| Significance based on thresholds: p Value: 0.050; r Value: 0.5; Fold change: 1.5 |  |  |
| Analysis Type | Significant Analyses | Total Unique Analyses |
| GFR Analyses | 6 | 43 |
| Serum Creatinine Analyses | 3 | 47 |
| Age Analyses | 2 | 44 |
| Disease vs. Control Analyses | 2 | 41 |
| Proteinuria Analyses | 2 | 46 |
| Transplant Analyses | 2 | 11 |
| Blood Pressure Analyses | 1 | 23 |
| Body Mass Index Analyses | 1 | 25 |
| BUN Analyses | 1 | 5 |
| Fasting Blood Glucose Analyses | 1 | 4 |
| Interstitial Fibrosis and Tubular Atrophy Analyses | 1 | 2 |
| ACR Analyses | 0 | 4 |
| APOL1 Risk Status Analyses | 0 | 7 |
| Disease vs. Other Disease(s) Analyses | 0 | 51 |
| Donor Type Analyses | 0 | 2 |
| Hemoglobin Analyses | 0 | 0 |
| Mesangial Hypercellularity Analyses | 0 | 1 |
| Post-Event Treatment Analyses | 0 | 1 |
| Race Ethnicity Analyses | 0 | 30 |
| Schena Grade Analyses | 0 | 1 |
| Segmental Glomerulosclerosis Analyses | 0 | 1 |
| Sex Analyses | 0 | 46 |
| Tissue Type Analyses | 0 | 34 |
| Weight Analyses | 0 | 32 |
| WHO Lupus Nephritis Class Analyses | 0 | 6 |
| Total | 22 | 507 |

Data obtained from NephroSeq v.5.0
www.nephroseq.org, September 2022, University of Michigan, Ann Arbor, MI

**Supplementary Table 2.**

| Data Type | Dataset | Analysis | Analysis Synopsis | Analysis Type | p-Value | Fold Change | r Value | Reporter |
| --- | --- | --- | --- | --- | --- | --- | --- | --- |
| mRNA | Nakagawa CKD Kidney | Chronic Kidney Disease vs. Normal Kidney (Discovery Set) | Disease vs. Control Analysis | over expression | 4.60E-06 | 1.807 |  | A_23_P109133 |
| mRNA | Flechner Transplant Blood | Renal Dysfunction vs. No Rejection (Cadaveric Donors) | Transplant Analysis | under expression | 1.30E-04 | -1.576 |  | 34020_at |
| mRNA | Woroniecka Diabetes TubInt | (All Measured Samples) | GFR (MDRD) Analysis | correlation | 0.003 |  | 0.598 | 207848_at |
| mRNA | Hodgin FSGS Glom | (Focal Segmental Glomerulosclerosis Samples) | Interstitial Fibrosis and Tubular Atrophy Analysis | correlation | 0.003 |  | 0.885 | g13259532_3p_at |
| mRNA | Reich IgAN TubInt | (IgA Nephropathy Samples) | Proteinuria Analysis | correlation | 0.003 |  | -0.623 | 207848_at |
| mRNA | Flechner Transplant Blood | (Cadaveric Donors) | GFR (MDRD) Analysis | correlation | 0.004 |  | 0.679 | 34020_at |
| mRNA | ERCB Nephrotic Syndrome TubInt | (Focal Segmental Glomerulosclerosis Samples) | GFR (MDRD) Analysis | correlation | 0.005 |  | 0.66 | ENSG00000101200 |
| mRNA | Hodgin FSGS Glom | (Focal Segmental Glomerulosclerosis Samples) | Serum Creatinine Analysis | correlation | 0.006 |  | 0.863 | g13259532_3p_at |
| mRNA | Gunther Transplant Blood | (Acute Rejection Samples) | Age Analysis | correlation | 0.007 |  | 0.583 | 207848_at |
| mRNA | ERCB Lupus TubInt | (Lupus Nephritis Samples) | BUN Analysis | correlation | 0.008 |  | -0.886 | ENSG00000101200 |
| mRNA | Kurian Transplant Kidney | Laparoscopic Donor Nephrectomy vs. Open Donor Nephrectomy (All Measured Samples) | Transplant Analysis | over expression | 0.008 | 2.803 |  | 207848_at |
| mRNA | Flechner Transplant Blood | (Cadaveric Donors) | Serum Creatinine Analysis | correlation | 0.01 |  | -0.621 | 34020_at |
| mRNA | Cox IgAN Blood | (IgA Nephropathy Samples) | GFR (CG) Analysis | correlation | 0.017 |  | 0.801 | ILMN_1811443 |
| mRNA | Sampson Nephrotic Syndrome TubInt | (Minimal Change Disease Samples) | Proteinuria Analysis | correlation | 0.026 |  | -0.665 | 551 |
| mRNA | Cox IgAN Blood 2 | IgA Nephropathy vs. Healthy Living Donor | Disease vs. Control Analysis | over expression | 0.037 | 1.739 |  | 207848_at |
| mRNA | Ju CKD Glom | (Minimal Change Disease Samples) | Serum Creatinine Analysis | correlation | 0.037 |  | -0.605 | 551 |
| mRNA | ERCB Lupus TubInt | (Lupus Nephritis Samples) | GFR (CG) Analysis | correlation | 0.038 |  | 0.782 | ENSG00000101200 |
| mRNA | Flechner Transplant Blood | (Cadaveric Donors) | Age Analysis | correlation | 0.042 |  | 0.513 | 34020_at |
| mRNA | Berthier Lupus TubInt | (Healthy Living Donors) | Blood Pressure Analysis | correlation | 0.046 |  | -0.997 | 207848_at |
| mRNA | Hodgin Diabetes Mouse Glom | (Diabetic Nephropathy Mouse Model eNOS-deficient C57BLKS db/db) | Fasting Blood Glucose Analysis | correlation | 0.046 |  | 0.762 | 11998 |
| mRNA | ERCB Nephrotic Syndrome TubInt | (Focal Segmental Glomerulosclerosis Samples) | GFR (CKD-EPI) Analysis | correlation | 0.046 |  | 0.641 | ENSG00000101200 |
| mRNA | Ju CKD Glom | (IgA Nephropathy Samples) | Body Mass Index Analysis | correlation | 0.049 |  | 0.516 | 551 |

Data obtained from NephroSeq v.5.0 www.nephroseq.org, September 2022, University of Michigan, Ann Arbor, MI

## Notes

https://www.ncbi.nlm.nih.gov/geo/query/acc.cgi?acc=GSE76632

https://www.ncbi.nlm.nih.gov/geo/query/acc.cgi?acc=GSE66494

### Competing Interest Statement

The authors have declared no competing interest.

### Summary of Updates

Updated supplemental images with a new transgenic mouse model and updated supplemental tables with information regarding gene expression.

